# Bacterial deamidases modulate ubiquitin structure and dynamics to dysregulate ubiquitin signaling

**DOI:** 10.1101/2023.05.22.541748

**Authors:** Rashmi Agrata, Priyesh Mohanty, Aravind Ravichandran, Sanju Kumari, Nishant Varshney, GS Arun, Kanchan Chauhan, Jess Li, Kalyan S. Chakrabarti, R. Andrew Byrd, Ranabir Das

## Abstract

The deamidases secreted by *Burkholderia pseudomallei* and *Enteropathogenic Ecoli* modify the residue 40 in ubiquitin from a Glutamine (Q) to Glutamate (E), triggering several downstream processes to cause cell cycle arrest and activate immune responses. Deamidation hampers the activity of ubiquitin and its interaction with ubiquitin chain receptors by an unknown mechanism. Here, we study the effect of deamidation on ubiquitin structure and dynamics. We report the crystal structure of the deamidated ubiquitin, supported by NMR and molecular dynamics simulations. The structure reveals a new intra-molecular salt bridge between the deamidated region and the C-terminal tail of ubiquitin. The salt bridge perturbs the dynamics of the ubiquitin tail to reduce affinity for ubiquitin receptors like the p62 ubiquitin-associated domain. The salt bridge disrupts the transition to catalytically active E2~Ub closed conformation. Consequently, RING E3s fail to interact with E2~Ub, reducing ubiquitination activity. Our findings reveal that deamidation-induced intramolecular salt bridges in ubiquitin modulate conformational ensembles to inactivate ubiquitination.

## Introduction

In eukaryotes, posttranslational modification (PTMs) enables proteome diversity essential for cellular processes^1^. Ubiquitination is a modification that regulates proteins’ degradation, molecular trafficking, interactions, and DNA repair, among others^2^. Ubiquitin, an 8.5 kDa small modifier, gets covalently attached to a substrate’s lysine residue with the sequential activity of three enzymes, E1, E2, and E3^3^. Activated by ATP and E1, the c-terminal tail of ubiquitin gets transferred onto E2 to form the E2~Ub^4, 5^. The thioester-linked E2~Ub is a dynamic ensemble of various conformations attributed to the flexibility of the ubiquitin tail^6^. E3 ligases bind to the E2~Ub intermediate to bring them in a catalytically closed conformation that aids nucleophilic attack by the ɛ-amino group of a lysine side chain from a substrate protein and thus allows isopeptide-linked attachment of ubiquitin onto the substrate^7^. Different E2/E3 pairs dictate the linkage specificity of ubiquitin chain^8^. These chains, varying in size, conformation, and linkages, regulate the fate of the substrate and thus control a wide range of cellular events^3, 8, 9^.

As the ubiquitination pathway plays a crucial regulator in diverse functions, it is a suitable target for pathogens to hijack a host cell for survival^10^. Bacterial pathogens inject a bunch of effector proteins upon infection using their T3SS/T4SS and have been shown to function as kinases, cycle-inhibiting factors (CIFs), E3-ligases, and E40UBs that interfere with the host ubiquitination^11, 12^. The recent emergence of glutamine deamidase, a *cif*-family effector from several gram-negative bacterial pathogens, highlighted a novel cross-talk between ubiquitin-like pathways and deamidation^13^. *Cif* from enteropathogenic *Escherichia coli* (EPEC) converts glutamine, Q40 in NEDD8 to a glutamate E40 via deamidation^14^. In contrast, a *Cif*-homolog from *Burkholderia pseudomallei* (CHBP) deamidates the same residue in both ubiquitin and NEDD8^15^ and leads to the cell-cycle arrest or actin-stress fiber formations in the host cells^16^. Cifs block the degradation of cell cycle regulators like p21/27, inactivating cycle-dependent kinases^17^. Moreover, CHBP inhibits TNF-α stimulated NF-ĸB activity by blocking the ubiquitin-mediated degradation of its inhibitor IĸBα^15^. Cellular phenotypes associated with the activity of the deamidases have been well-characterized. However, the structural changes in ubiquitin and the molecular mechanism underlying the dysfunction of the ubiquitin pathway are poorly understood. *In vitro* ubiquitin chain synthesis reduces after CHBP treatment^15^, suggesting a possible molecular mechanism that inactivates ubiquitination.

In this study, we investigate the changes in ubiquitin and ubiquitination activity by employing biophysical techniques, MD simulations, and biochemical assays. We report that deamidation of ubiquitin (E40Ub) creates a new intra-molecular salt bridge between positively charged arginine R72/74 in the c-terminal tail and the negatively charged glutamate E40, introduced upon deamidation, without perturbing the globular ubiquitin fold. Our study also suggested that deamidation hampers ubiquitin activation by E1, leading to reduced E2~Ub conjugation. Molecular dynamics simulations revealed an altered conformation of E2~E40Ub that destabilizes the catalytically closed conformation, validated using NMR relaxation techniques. We further examined the dynamics, interactions, and activity of ubiquitin in the E2~Ub conjugate and found that deamidation-induced conformational change hinders interactions with RING E3s leading to reduced ubiquitination. Our study provides a detailed mechanism wherein a pathogen adversely modulates ubiquitin activity by introducing a negative charge in ubiquitin to hijack the host cell machinery.

## Materials and Methods

### Molecular dynamics simulations

The crystal structure of E40Ub reported in this study was used as a starting structure for E40Ub simulations. The E1-Ub complex was taken from PDB 6DC6. The closed conformation of Ubc13~Ub and the Ubc13~Ub-RNF4^RING^ complex was extracted from PDB 5AIU and 5AIT, respectively. The extended conformation of Ubc13~Ub was modeled by the superposition of Ubc13 (PDB 3HCU) onto the extended conformation of UBCH5B~Ub (PDB 3JVZ). Glutamine-to-glutamate substitutions were introduced by replacing existing sidechains with best-aligning rotamers from the Dunbrack rotamer library using the rotamer tool available in UCSF Chimera. Electrostatic surface potentials were calculated using the Adaptive Poisson-Boltzmann Solver.

Conventional MD simulations were performed in GROMACS 4.6, and SMD simulations were performed using the pull code in GROMACS 5.1.2. The simulation system for E40Ub was modeled using the AMBER99SB*-ILDN force field, while the AMBER99SBws force field was used for all other systems to prevent overestimating protein-protein interactions. Na^+^ and Cl^−^ ions were modeled using parameters derived by Joung and Cheatham. Non-bonded corrections for cation-chloride/carboxylate and amine-carboxylate interactions derived by Yoo and Aksementiev were applied for both force fields. AMBER99 compatible parameters for the thioester bond in Ubc13~Ub and zinc coordination sites in RNF4^RING^ were adopted from previous studies.

The starting structures for conventional MD runs of E40Ub and Ubc13~Ub/E40Ub were solvated in a rhombic dodecahedron box by adding TIP3P/TIP4P2005 water molecules and 0.15 M NaCl. A suitable number of counter ions were added to neutralize the residual charge of the system. A minimum distance of 1.4 nm between any solute atom and the box edge was specified. SMD simulations of the E1-Ub complex were performed in a cuboidal box of dimensions: 15.0 (x) nm x 13.0 (y) nm x 13.0 (z) nm. Dissociation of Ubc13~Ub/Ubc13~Ub-RNF4^RING^ complexes was performed in a cuboidal box of dimensions: 12.6 (x) nm x 9.0 (y) nm x 9.0 (z) nm.

The electrically neutral, solvated system was then subjected to energy minimization using the steepest descent method for a maximum of 5000 steps until the maximum force on any atom was less than 1000 kJ mol^−1^ nm^−1^. Temperature and pressure equilibration was performed for 500 ps using the v-rescaling thermostat (τ_t_ = 0.1 ps) and Berendsen barostat (τ_p_ = 1.0). Numerical integration was performed using the leapfrog integrator using a time step of 2 fs. All bond lengths were constrained with the LINCS algorithm, and harmonic restraints (k = 1000 kJ mol^−1^ nm^−2^) were applied to all heavy atoms of proteins to prevent structural distortion during equilibration.

Production simulations were performed under periodic boundary conditions at 300 K and 1 bar pressure (NPT ensemble). In addition, virtual interaction sites were employed for hydrogen atoms, which permitted a 5 fs time step. Temperature and pressure control were achieved using the v-rescaling thermostat (τ_t_ = 2.5 ps) and Parrinello-Rahman barostat (τ_p_ = 5 ps). Short-range electrostatics and van der Waals interactions were calculated using a 1.0 nm cutoff. Long-range electrostatics were computed using the PME method. Long-range van der Waal’s interactions were treated using an analytical dispersion correction. SMD simulations of E1-Ub/Ubc13~Ub/Ubc13~Ub-RNF4^RING^ were performed for 20 ns at a pull rate of 0.25 nm ns^−1^. The pull force was applied on the COM of Ub (aa:1-70) through a dummy atom attached to it via a harmonic spring with a force constant (k) of 1500 kJ mol^−1^ nm^−2^. The COM motion of E1/Ubc13 was removed every 100 fs to promote the unbinding of Ub.

For analysis of conventional MD runs, system coordinates were stored every 0.25 ns for subsequent analysis. For the analysis of SMD runs, forces, and COM distances were calculated every 5 ps. Salt-bridge occupancies were computed using a cutoff distance of 0.5 nm between Cζ (Arg) and Cγ/δ (Asp/Glu) atoms. Minimum distances between Ubc13 and Ub (aa: 1-70) were calculated using the *g_mindist* tool in GROMACS. The coulombic interaction energies were calculated using *g_energy*. *g_sgangle* was used to calculate the angle between the pre-defined vectors for the helices of Ubc13 and Ub. RMSD-based structural clustering was performed using the GROMOS algorithm. Briefly, RMSD matrices were calculated for the concatenated, 1 μs macrotrajectories of Wt and mutant conjugates, after which an RMSD cutoff of 0.55 nm was applied to similar E2~Ub conformations and identify representative structures for each cluster. The RMSD was calculated for the entire Ubc13~Ub backbone.

### Targeted Molecular Dynamics simulations

Systems for TMD were prepared with the LEaP program of Ambertools18. Systems were prepared in a cubic box of TIP3P water, with a minimum distance of at least 1.2 nm between solute atoms and the box edge. Counter ions were added to neutralize each system. Parameters describing system topology were based on the Amberff99SB-ildn force field. The distance cutoff for short-range non-bonded interactions was set to 1 nm. The particle mesh Ewald (PME) method was employed to treat long-range electrostatic interactions. The SHAKE algorithm constrained all bonds involving hydrogen atoms. The temperature was set to 300 K using a Langevin thermostat with a collision frequency of 1.0 ps^−1^. The Pressure was maintained at 1 bar using the Berendsen barostat with a coupling constant of 1.0 ps. Using the hydrogen mass repartitioning (HMR) scheme, the integration time step was set to 4 fs. Dynamics were propagated using the leapfrog integrator.

The systems were first relaxed by energy minimization using the Sander module of Amber18. The respective systems were then heated from 0 K to 300K for 200 ps (NVT equilibration), during which positional restraints (10 kcal/mol/Å^2^) were applied to all protein atoms. Subsequently, a 400 ps NPT MD was run to adjust the system’s density to 1 g/cm^3^. In the production run, TMD was used to drive the system from open to closed conformation for both Ub and E40Ub conjugates. The backbone atoms corresponding to Ubc13 (initial conformation) were used for superposition, and Ub backbone atoms (aa: 1-70) were used for calculating RMSD. In the TMD cycle, T_RMSD_ was reduced by 0.5 nm every 200 ps, followed by 40 ps of equilibration. The cycle was repeated until the T_RMSD_ reached 0 Å. TMDs were performed at five k values (0.01, 0.005, 0.001, 0.0005 & 0.0001). A harmonic restraint of 100 kcal/mol was applied between Glu40 (Oε) and Arg72 (Nη) atoms to keep the salt bridge intact. At each force constant, five TMD replicates were performed.

### Cloning and mutagenesis

Ubiquitin mutants (Ub Q40E, Q40A, S20C, R72A, D39A) were cloned using PCR-based site-directed mutagenesis with no affinity tag. His-Ubc13 in the pET24 vector, GST-Ubc13 C87K/K92T/K94Q in the pGEX6P1 vector, and GST-RNF38^RING^ (residues 387–465) in the pGEX4T1/UbE2D2 S22R C85K in RSF Duet vector were a gift from Prof. Rachel Klevit, Prof. Catherine Day and Prof. Danny Huang respectively. All the plasmid sequences were confirmed by sanger sequencing.

### Protein expression and purification

All proteins were expressed in BL21 DE3 star cells (Invitrogen), grown at 37°C (in LB media for unlabelled proteins and M9 media for ^13^C or ^15^N isotope-labeled proteins) till OD_600_ reached 0.7, and were induced with 0.5 mM IPTG for 4–5 hr. The harvested cells were lysed in 50 mM Tris-HCl, 250 mM NaCl, 5 mM βME, 2% glycerol, 0.01% triton-X, 1 mM PMSF, and DNase at pH 8 using Emulsiflex high-pressure homogenizer. All the GST-tagged proteins were bound to GSTrap HP columns (GE Healthcare) and were eluted with 10 mM reduced glutathione. All the His-tagged proteins were bound to HisTrap HP columns (GE Healthcare) and were eluted with 100 mM-500 mM Imidazole gradients. To purify Ub and its variants, lysed cells were resuspended in 50 mM Sodium acetate, 5 mM BME, 0.01% triton-X, pH 4.5 by sonication, and centrifuged to pellet cell debris. The supernatant was passed through the SP FF column (GE Healthcare), and the protein was eluted by gradient elution of increasing salt concentration (0 mM NaCl-600 mM NaCl). The proteins were further purified by gel filtration on Superdex 75 pg 16/600 (GE Healthcare) in 25 mM Tris-HCl, 100 mM NaCl and 2 mM βME at pH 7.5. For Alexa labeling Ub/E40Ub, the S20C mutant was purified and incubated with excess Alexa Fluor^TM^ 488 C_5_ Maleimide (Invitrogen: A10254) at 4°C overnight. Free Alexa was removed using a desalting column, and the Alexa-tagged proteins were stored.

### E1-Ub binding assay

E1-Ub binding studies were performed using fluorescence anisotropy measurements and NMR. Samples containing complexes of serially diluted E1 from 3 µM to 1.4 nM in 25 mM Tris-HCl, 50 mM NaCl (pH 7.5), and 5 nM Alexa-labelled Ub/E40Ub were prepared in 96-well flat-bottom black non-binding polystyrol plate (Coster). Fluorescence polarisation was measured at λ_ex_=470 nm and λ_em_=516 nm in the Tecan Infinite M1000 PRO microplate reader at 25°C, and the raw parallel and perpendicular intensities were used to determine the changes in anisotropy upon E1 binding. Kd was calculated by fitting the curve in Sigmaplot. To monitor chemical shift perturbations (CSPs) and change in intensities of Ub/E40Ub peaks upon E1 binding, 180 µL of equimolar (65 µM) ^15^N/D_2_O labeled Ub/E40Ub, and E1 was prepared in 25 mM Tris-HCl, 50 mM NaCl (pH 7.5), 10% D_2_O. TROSY spectra were collected in an 800 MHz Bruker Avance III HD spectrometer AT 298K.

### E1-Ub and E2-Ub activation assays

To monitor the E1-Ub activation rate, 0.5 µM E1 and 500 µM ^15^N labeled Ub mutants were incubated with 50 mM Na_2_HPO_4_, 100 mM MESNA (Sigma: M1511), 10 mM ATP, and 10 mM MgCl_2_ (pH 6.7) at 30°C. An equal volume of aliquots (20 µL) was taken at 0, 0.5, 1, 2, and 4 hours and the reaction was quenched with 20 mM EDTA. 500 µM Ub mutants (unlabeled) were added for the internal control, and the reactions were dialyzed in water and the samples were prepared by mixing 0.4%(v/v) TFA. The samples were injected into the ESI-MS in triplicate, and the data were analyzed using ThermoBiopharmaFinder.

The rate of Ub/E40Ub discharge from E1~Ub to E2 (Ubc13) was monitored using a trans-thiolation assay. 0.5 µM E1 and 1 µM Alexa-labelled Ub/E40Ub were incubated in 25 mM Tris-HCl, 50 mM NaCl (pH 7.5), 2.5 mM ATP, and 5 mM MgCl_2_ at 30°C for 10 min. The reaction was quenched using 200 mM EDTA, and the discharge was started with 1 µM Ubc13 at 16°C. Equal aliquots were taken out at 0, 0.3, 1, 2, and 5 mins and loaded onto the 15%-SDS gel. The images were taken in Amersham Typhoon^TM^ Imager (GE) and quantified using ImageJ.

To monitor E2~Ub formation, active site Cys-Ser mutants were generated for Ubc13, UbcH5B, and UbE2B. 0.5 µM E1, 5 µM E2s, and 20 µM Ub/E40Ub were mixed in 25 mM Tris-HCl, 100 mM NaCl, 5 mM ATP, and 5 mM MgCl_2_ at pH 7.5. Equal aliquots of reaction mixes were taken out at different time points, loaded onto the 15%-SDS gel, and stained with SYPRO Ruby red. The images were taken in ImageQuant LAS4000 and quantified using ImageJ.

### Ubc13~Ub isopeptide conjugate purification

Ubc13 active site Cys-Lys mutant C87K/K92T/K94Q generated the stable isopeptide-linked Ubc13~Ub/E40Ub conjugates. 0.5 µM E1, 50 µM Ubc13, and 100 µM Ub/E40Ub were incubated in 25 mM Tris-HCl (pH 8.5), 10 mM phosphocreatine (Sigma), 1 mM ATP (Sigma), 5 mM MgCl_2_ and 1U/ml creatine phosphokinase (Sigma) at 37°C for 25-30 hours. The reaction mixture was loaded onto GE mono Q^TM^ 10/100 GL with 25 mM Tris-HCl (pH 8.0), 10 mM NaCl and 1 mM BME, and the conjugates were eluted with a salt gradient of 0-400 mM NaCl in the loading buffer. Ubc13~Ub/E40Ub peak fractions were run on 15% SDS-gel, and pure fractions were concentrated to make 0.5-4 mg/mL samples.

### Analytical gel filtration

Superdex 75 10/300 GL (GE Healthcare) was equilibrated with 25 mM Tris-HCl, 100 mM NaCl, and 2 mM βME (pH 7.5). Samples of 0.6 mg/mL of Ubc13 and Ubc13~Ub/E40Ub were injected with a constant 0.4 mL/min flow rate at 4°C. Protein analytical standard solution (Sigma) was used for plotting the standard chart for Stoke’s radius determination.

### NMR titrations

The NMR samples were prepared in 25 mM Tris, 100 mM NaCl, pH 7.5, and 10% D_2_O. For titration, ~ 1.6 mM RNF38^RING^ were titrated into 0.150 mM ^15^NUbc13, 0.18 mM Ubc13-^15^NUb and 0.15 mM Ubc13-^15^NE40Ub. The titration data were fit to a 1:1 protein:ligand model using the equation CSP_obs_ = CSP_max_ {([P]_t_+[L]_t_+K_d_) − [([P]_t_+[L]_t_+K_d_)^2^-4[P]_t_[L]_t_]^1/2^}/2[P]_t_, where [P]_t_ and [L]_t_ are total concentrations of protein and ligand at any titration point. To titrate RNF38^RING^ to Ube2d2, 1.4 mM of RNF38^RING^ was titrated to 0.125 mM Ube2d2, 0.12 mM of Ube2d2~Q40Ub, and 0.12 mM of Ube2d2~E40Ub. For p62-UBA/Ub titrations, 3 mM of the p62-UBA domain was titrated to 0.2 mM ^15^N-labeled Q40Ub or E40Ub. All NMR experiments were recorded at 298K on 600 MHz or 800 MHz Bruker Avance III HD spectrometer.

### Relaxation dispersion experiments and analysis

The high-power relaxation experiments^32^ were measured using the constant time CPMG experiment for quantifying the micro-to-millisecond timescale exchange process in Q40Ub and E40Ub. All experiments were carried out in Bruker Avance 600 MHz and 800 MHz spectrometers fitted with cryoprobe at 277 K. The constant time (CT) CPMG delay is divided into two halves, sandwiching the U-element that ensures the equal contribution of anti-phase and in-phase relaxation to R_2,eff_ in all frequencies. The refocusing pulses are applied with strong B_1_ fields for all refocusing frequencies, thus reducing any off-resonance effects in R_2,eff_ measurement. The R_2,eff_ values were measured at CPMG frequencies (ν_CPMG_) of 66.7 Hz, 133 Hz, 267 Hz, 400 Hz, 533 Hz, 667 Hz, 1333 Hz, 2000 Hz, 256 Hz, 3333 Hz, 4000 Hz, 4667 Hz, 5333 Hz, and 6000 Hz. The constant volume of 200 μL of NMR samples was put inside 3 mm NMR tubes in 25 mM Tris, 100 mM NaCl, pH 7.5 buffer, and 10% D2O. In all experiments, the ^15^N-ubiquitin concentration was 1 mM. The p62-UBA (unlabeled) concentration was changed from 0, 0.02 mM, 0.05 mM, 0.1 mM, 0.25 mM up to 0.5 mM. The R_2,eff_ was calculated as R_2,eff_ (ν_CPMG_) = −log (I(ν_CPMG_)/I_0_)/T, where ν_CPMG_ is the effective frequency of the CPMG field, T is the constant delay (60 ms), I_0_ is the intensity of the peak in reference experiment and I(ν_CPMG_) is the intensity of the peak at that particular CPMG frequency. The experiment was performed with 3 sec recycle delay between increments using 12 different refocusing field strengths between 0 to 6000 Hz collected in a scrambled and interleaved manner with 1024 (^1^H) and 130 (^15^N) complex points, respectively. There is a heat compensation block in the middle of the recycle delay to dump the extra CPMG cycles so that the total number of CPMG 180 refocusing pulses at fixed B_1_ field strength is identical during the individual scans. The global uncertainty for the experimental data was calculated from the average standard deviation of R_2,eff_ values for 5 residues that show flat dispersion profiles. The residue-specific uncertainties were calculated by measuring the deviation between repetitions of identical frequency (667 Hz) of the R_2,eff_ values. The largest value between the global or residue-specific uncertainties is reported. The ligand-free experiment was repeated for the E40Ub, with all other parameters unchanged.

We fitted the relaxation rates R_2,eff_ with the fast-exchange formula

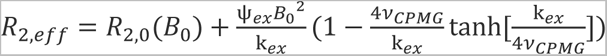

The two NMR data sets for R_2,eff_ as a function of *ν_CPMG_* - at the two ^15^N resonance frequencies, 60 MHz and 80 MHz were jointly fitted using the four fit parameters R_2,0_(60 MHz), R_2,0_ (81 MHz), ψ_ex,_ and k_ex_ to determine the kinetics experienced by the ^15^N labeled ubiquitin. We used the function NonlinearModelFit of Mathematica 11.3 in these fits. The fits were 1/ ΔR ^2^ weighted, and the errors of the fit parameters were estimated from the fit residuals with the standard variance estimator function of NonlinearModelFit.

### Protein Crystallization, diffraction, and data collection

For crystallization, 15 mg/ml E40Ub samples were grown in 0.1M Sodium Acetate Trihydrate pH 4.5, 25%PEG 3350 at 25°C. The hanging drops were set up by mixing 1:1 protein and precipitant solutions. The crystals diffracted without any cryo-protectant. The crystals were frozen at 100K directly on a stream of liquid nitrogen from the cryo-probe. Data were recorded in-house on an R-AXIS IV++ image plate at 1.5148 Å using Cu-Kalpha radiation generated by FR-X from Rigaku Inc. The crystal-to-detector distance was kept at 100mm. Successive frames were collected using a crystal oscillation of 1.0°. An expert pipeline *XIA2* (winter *et al*., 2013) from *CCP4i2* (Potterton *et al.*, 2018) was used to automate data processing from diffraction images to scaled and merged data. The diffracted data were indexed and integrated using *XDS* (Kabsch, 2010) and scaled using *AIMLESS* (Evans & Murshudov, 2013). Final data reduction was performed using *AIMLESS* in *CCP4i v7.0.072* (Potterton *et al.*, 2002), taking unmerged and unscaled data for better statistics. 5% of the data was reserved for cross-validation. Data collection statistics are provided in Table 1.

### Structure determination and refinement

The crystal structure of wild-type ubiquitin (PDB ID: 1UBQ) was used as the template for the search model. The structure was determined using the molecular replacement method using data between 35.24 and 1.6 Å in the PHASER (McCoy et al., 2007) from the PHENIX suite (Adams et al., 2010). Initial model building was performed using *AUTOBUILD* (Terwilliger *et al*., 2008), and initial structure refinement was performed using *PHENIX.REFINE* (Afonine *et al*., 2012). Manual corrections and fitting based on observed electron density were done using *COOT* (Emsley *et al*., 2010). Final refinement using the maximum likelihood method was carried out by multiple cycles in *REFMAC5* (Murshudov *et al.*, 2011) in *CCP4i v7.0.072*. Water molecules were manually added in coot at peaks of density above 4σ in the asymmetric unit. The model’s geometry was checked using *MolProbity* (Chen *et al.,* 2010). The structure is submitted to the protein data bank (PDB ID: 7F0N).

## Results

### Ubiquitin deamidation creates a new intra-molecular salt-bridge

Q40 deamidation of ubiquitin reduces the polyubiquitination for several E2 enzymes (Figures 1A, B, and S1A). The Q/E modification also creates a significant negatively charged surface patch in the nearby region, including the acidic residues D39 and E40 (Figure S1B, C). The structural changes in ubiquitin by deamidation were initially probed by NMR spectroscopy. Standard triple resonance experiments assigned the NMR chemical shifts of E40Ub atoms (Figure S1D). The chemical shifts of backbone amide resonances were similar between Q40Ub and E40Ub, suggesting subtle changes in the overall ubiquitin fold (Figure 1C). Chemical shift differences were detected around residue 40, expected as the deamidation site (Figure 1D). Interestingly, the C-terminal tail of ubiquitin showed significant chemical shift differences between Q40Ub and E40Ub. The C-terminal tail of ubiquitin includes two Arginine amino acids, R72 and R74. The sidechain Nε-Hε chemical shifts of R72 and R74 were altered in E40Ub, suggesting major structural changes in that region (Figure 1E).

**Figure 1.**
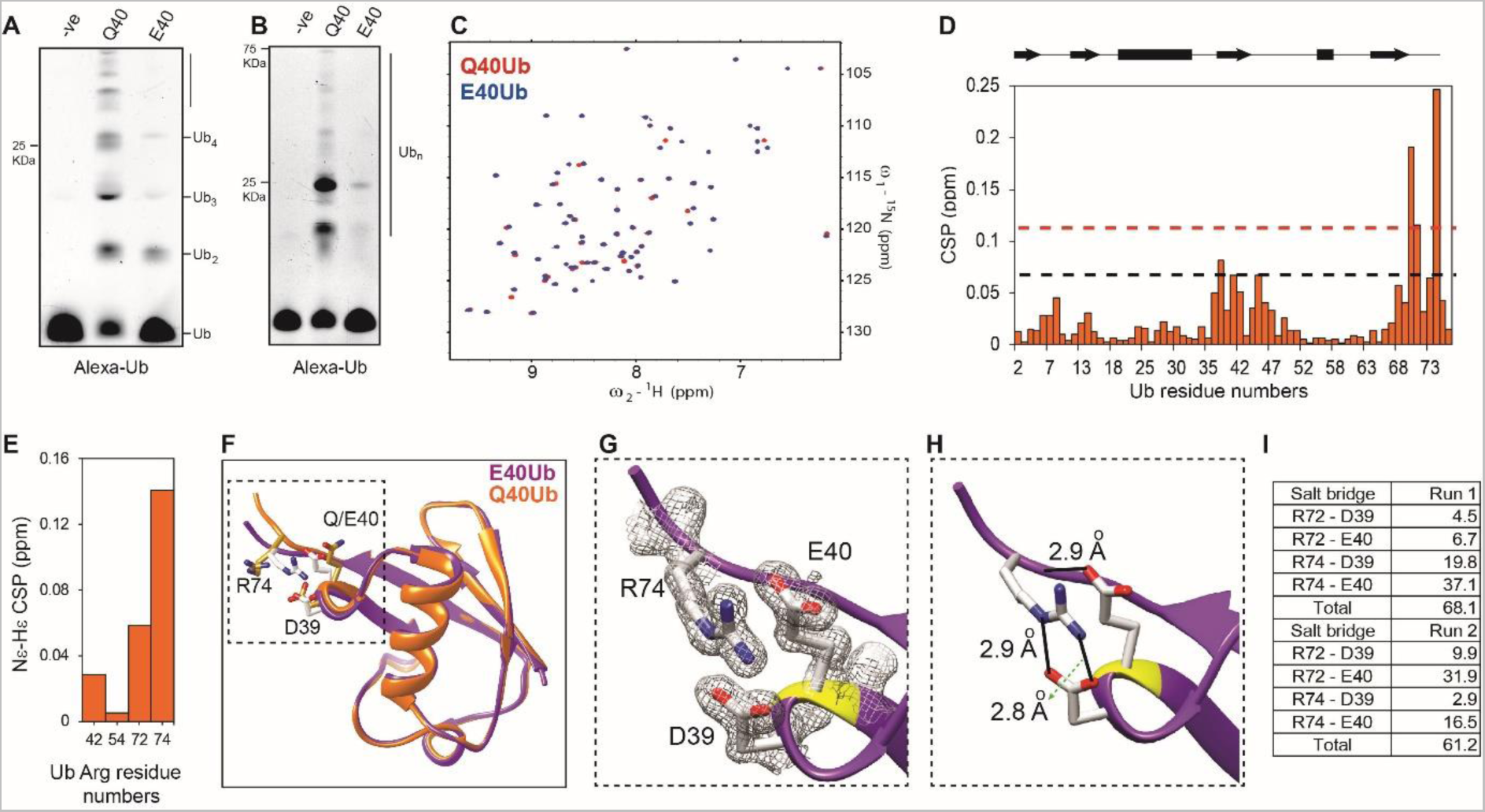
Deamidation induces the formation of new salt bridges in ubiquitin. (A) In-vitro ubiquitination reaction was performed using Ubc13 as the E2, RNF38^RING^ as the E3, and E40Ub or Q40Ub for 10 min. Mms2 was used as a co-factor in the reaction. The –ve lane is the same reaction without ATP. (B) The same assay as in (A) is repeated with the E2 Ubch5b. (C) An overlay of ^1^H-^15^N HSQCs of Q40Ub and E40Ub. (D) Chemical Shift Perturbations (CSPs) in amide resonances between Q40Ub and E40Ub are plotted against the Ub residue numbers. Residues in the c-terminal tail and around E40 show changes in the chemical environment. The secondary structure of Ub is shown above the plot. The broken black line is Standard Deviation (SD), and the red line is 2*SD. (E) CSPs plotted for Arginine Nε-Hε resonances between Q40Ub and E40Ub. (F) A superposition of crystal structures of Q40Ub (PDB id: 1UBQ) and E40Ub (this work). (G) A zoomed panel shows the electron density and packing of R74 between D39 and E40. (H) The new salt bridge formed between R74 and E40 is shown as black lines. A new hydrogen bond is also formed between E40 and the R72 backbone. The length of salt bridges and hydrogen bonds are mentioned. (I) Salt-bridge occupancies were calculated from two independent MD runs of E40Ub using a 0.5 nm cutoff.

E40Ub was crystallized to detect the structural changes upon deamidation. The crystal of E40Ub diffracted to 1.4 Å resolution, and the structure was determined using the wt ubiquitin as the model (Table S1). The Cα-RMSD between Q40Ub and E40Ub were within 1 Å for the ordered region, confirming no structural changes in the ordered regions (Figure 1F). However, the C-terminal end deviates from the conformation in the Q40Ub structure. The R74 side chain rotates and packs between D39 and E40 (Figure 1G). R74 sidechain Nε and Nη atoms form salt-bridges with D39 sidechain oxygen atoms. E40 forms a new hydrogen bond with the backbone of L73 (Figure 1H). All-atom molecular dynamics simulations studied the dynamics of these salt bridges, carried out for 0.5 μsec in E40-Ub in two separate runs. The simulations suggested the transient formation of multiple salt bridges between the C-terminal basic amino acids R72/R74 and the acidic patch D39/E40 (Figure 1I and S2).

### The salt bridge perturbs non-covalent interactions with the UBA domain

It is unclear how deamidation reduces the non-covalent interactions between ubiquitin and its receptors. The interaction of ubiquitin with the p62-UBA domain was measured by NMR titrations (Fig 2A, B). The dissociation constant of the complex between E40Ub and p62-UBA was three-fold lower than Q40Ub (Fig 2C, 2D). The structure of p62-UBA/Q40Ub was solved by NMR (Fig 2E, S3, and Table S2). The helices α1 and α3 in the UBA domain bind to the L8-I44-V70 hydrophobic patch in ubiquitin. Hydrophobic residues L8 and V70 in ubiquitin pack against the UBA domain’s hydrophobic residues M404 and L428 (Figure S3C). I44 interacts with Y433. Additionally, R74 and R72 form critical salt-bridge interactions with residue D429 in p62-UBA (Figure S3D). In E40Ub, the R74 and R72 are involved in intramolecular interactions within ubiquitin and are unavailable to form the necessary interaction with D429 in p62-UBA (Figure 2F). The loss of strong electrostatic interactions across the interface reduces the affinity of UBA/ubiquitin interactions.

**Figure 2.**
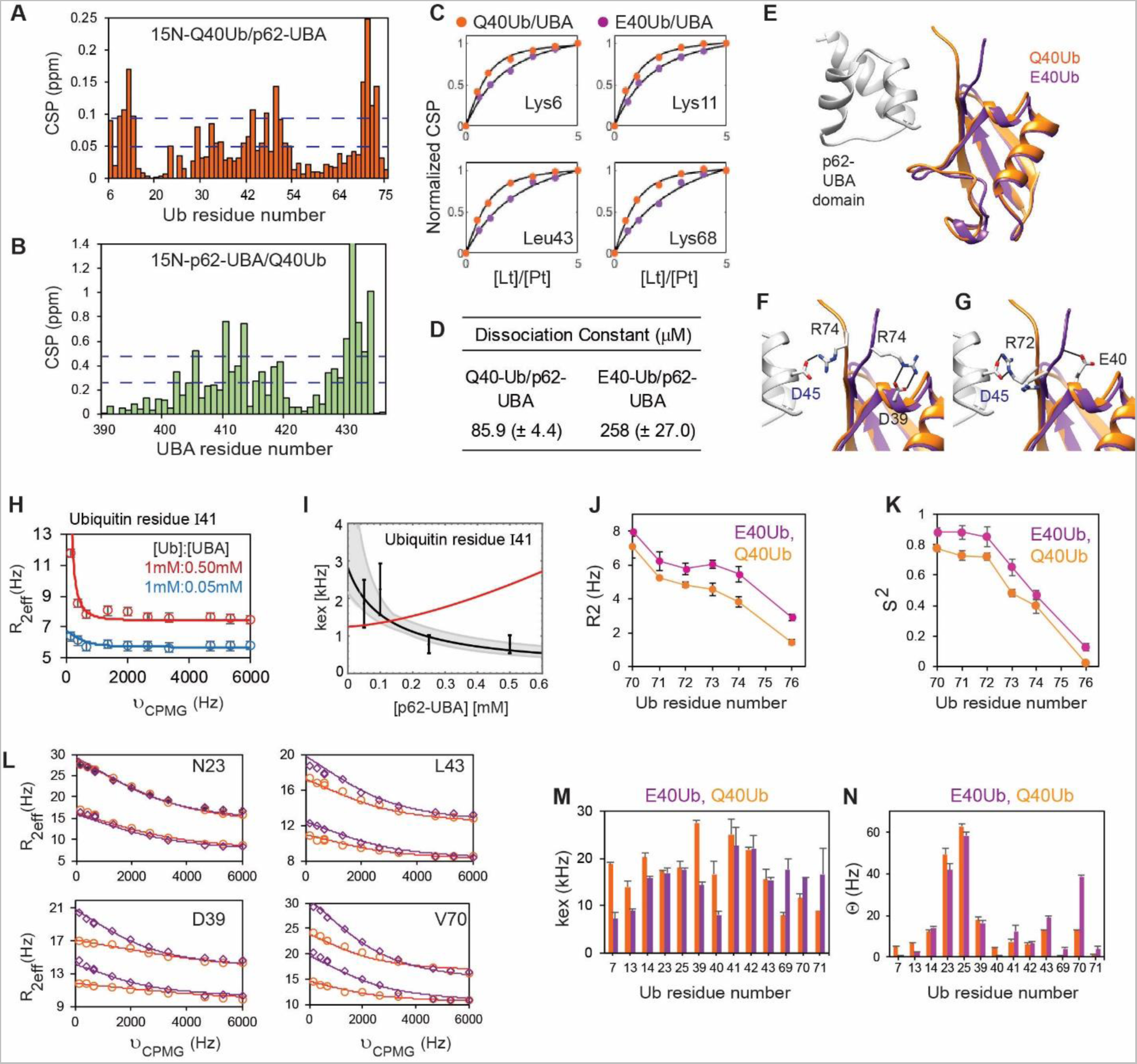
Deamidation perturbs the conformational dynamics and non-covalent interactions of Ub. (A) The chemical shift perturbation (CSP) in Ub, when bound to the p62-UBA domain. (B) The CSPs are plotted for the p62-UBA domain when bound to Ub. (C) The CSPs of Q40Ub and E40Ub when titrated with various concentrations of p62-UBA. (D) The dissociation constants of UBA/Ub complexes are obtained by fitting the titrations in (C). (E) The structure of the p62-UBA(white)/Q40Ub(orange) complex is shown. The E40Ub (purple) is superimposed on the Q40Ub structure. (F) The intermolecular salt bridge between D45 in UBA and R74 in Q40Ub is highlighted in black. The R74 in E40Ub is orientated in the opposite direction due to the intramolecular salt bridge with D39. (G) The intermolecular salt bridges between D45 in UBA and R72 in Q40Ub are highlighted. The C-terminal tail in E40Ub is distant from the UBA domain due to the hydrogen bond with E40. The R72 is orientated away from a potential intermolecular salt bridge with D45. (H) ^15^N relaxation dispersion for the amide of the ubiquitin residue I41 at 80 MHz in the presence of 0:05 mM and 0:5 mM of p62-UBA domain (blue and red data point). (I) The obtained exchange rates kex of the ubiquitin residue 41 (black data points with error bars) decrease with increasing p62-UBA concentration, indicating conformational selection. The gray lines with shaded error regions result from fits of the k_ex_ equations of the two-state and conformational-selection binding mechanism. The red line denotes the induced fit model. (J) and (K) are the order parameters (S^2^) and spin-spin relaxation rates (R_2_), respectively, of the C-terminal tail residues in Q40Ub and E40Ub. The same for complete protein is provided in Figure S4. (L) The ^15^N relaxation dispersion data (R_2eff_ against ν_CPMG_) are plotted for a few selected residues in Q40Ub (orange) and E40Ub (purple). (M) and (N) plots the k_ex_ and Θ (p_A_p_B_Δω^2^), respectively, for the residues with high R_ex_.

Ubiquitin binds the CIN85 SH3 domain by conformational selection, and the ubiquitin C-terminal region is critical for the selection^32^. The ubiquitin C-terminus is dynamic and adopts multiple conformations, where an extended conformation is exclusively selected for binding. To test whether ubiquitin binds to the p62-UBA domain by induced-fit mechanism or by conformational selection, we measured the concentration-dependent kinetics of Ub/p62-UBA interaction. Decreased exchange rate k_ex_ with increasing ligand p62-UBA concentration suggested ubiquitin binds the p62-UBA domain by conformational selection (Figure 2 and S4). Interestingly, both p62-UBA and ubiquitin form the complex by the conformational selection, which is distinct from SH3/ubiquitin interaction, where though ubiquitin binds by the conformational selection, SH3 binds by the induced-fit mechanism.

Given that C-terminal interactions and conformations are crucial for interaction with the p62-UBA domain, we tested whether the new salt bridge changes its dynamics to impact the interaction. The C-terminal’s fast backbone dynamics (ns-ps timescale) were compared in Q40- and E40Ub by standard R1, R2, and heteronuclear NOE experiments. The R2 and order parameter (S^2^) values increased significantly in E40Ub, indicating that deamidation makes the C-terminal more rigid (Figure 2J, K). To measure the changes in dynamics at the μs-ms timescales, we compared the relaxation dispersion profiles of Q40Ub and E40Ub. The high power relaxation dispersion experiments can pulse with frequency (ν_CPMG_) 6 kHz and measure dynamics up to 36 kHz (2πν_CPMG_). These experiments detect a peptide cis-trans flip motion in ubiquitin between E24 and G53, reflecting a relaxation dispersion profile for residues I23 and N25. The peptide flip motion is unaltered between the Q40 and E40Ub (Figure 2L). However, the motion of several residues at the deamidation site (D39, Q40, L43) changes in E40Ub. The exchange values (k_ex_) drop significantly at the deamidation site, suggesting that the salt bridge rigidifies the region (Figure 2M). Moreover, the Θ(p_A_.p_B_.(Δω)^2^) value between Q40Ub and E40Ub changes in residues 69 and 70, suggesting that the salt-bridge changes the conformational equilibria is the region. Overall, p62-UBA binds to Q40Ub through the conformational selection of a minor state, whose population decreases upon deamidation, reducing the binding constant of the interaction (Figure S5). The structural changes due to deamidation are local, but it has a broader impact on the ubiquitin dynamics and its noncovalent interactions.

### Ubiquitin deamidation modulates E1 activation

We then investigated the reduced ubiquitination activity upon deamidation. Ubiquitin has multiple covalent/non-covalent interactions during the ubiquitination reaction with the E1 and E2 enzymes, mediated by the hydrophobic patch and the C-terminal tail. It is unclear how deamidation affects these interactions and the catalytic rates of these enzymes. We first studied the interactions between ubiquitin and the Ub-activating enzyme E1. To probe the E1/ubiquitin non-covalent interaction, the ^15^N-TROSY HSQC spectra were collected for free Q40Ub and in the presence of equimolar E1 (Figure S6A, B). The peak intensities of backbone amide resonances reduced drastically for the regions around the hydrophobic patch (L8, I44, V70) as observed previously (Figure 3A)^20^, suggesting that ubiquitin non-covalently binds to E1 via the hydrophobic patch. However, E40Ub behaved similarly (Figure 3A), indicating no effect of deamidation on E1/ubiquitin binding. Similarly, the Chemical Shift Perturbations (CSPs) pattern was similar between Q40Ub and E40Ub (Figure 3B). To quantitatively study the effect of deamidation on E1/ubiquitin binding, we titrated fluorescently labeled Q40Ub with E1 and measured the changes in fluorescence anisotropy in the labeled ubiquitin. The change in anisotropy against ligand concentration was fit to extract the dissociation constant (K_d_). Both Q40Ub and E40Ub yielded similar K_d_ values of 72±6 nM and 80±7 nM, respectively, indicating that the strength of E1/ubiquitin binding is unaffected by deamidation (Figure 3C).

**Figure 3.**
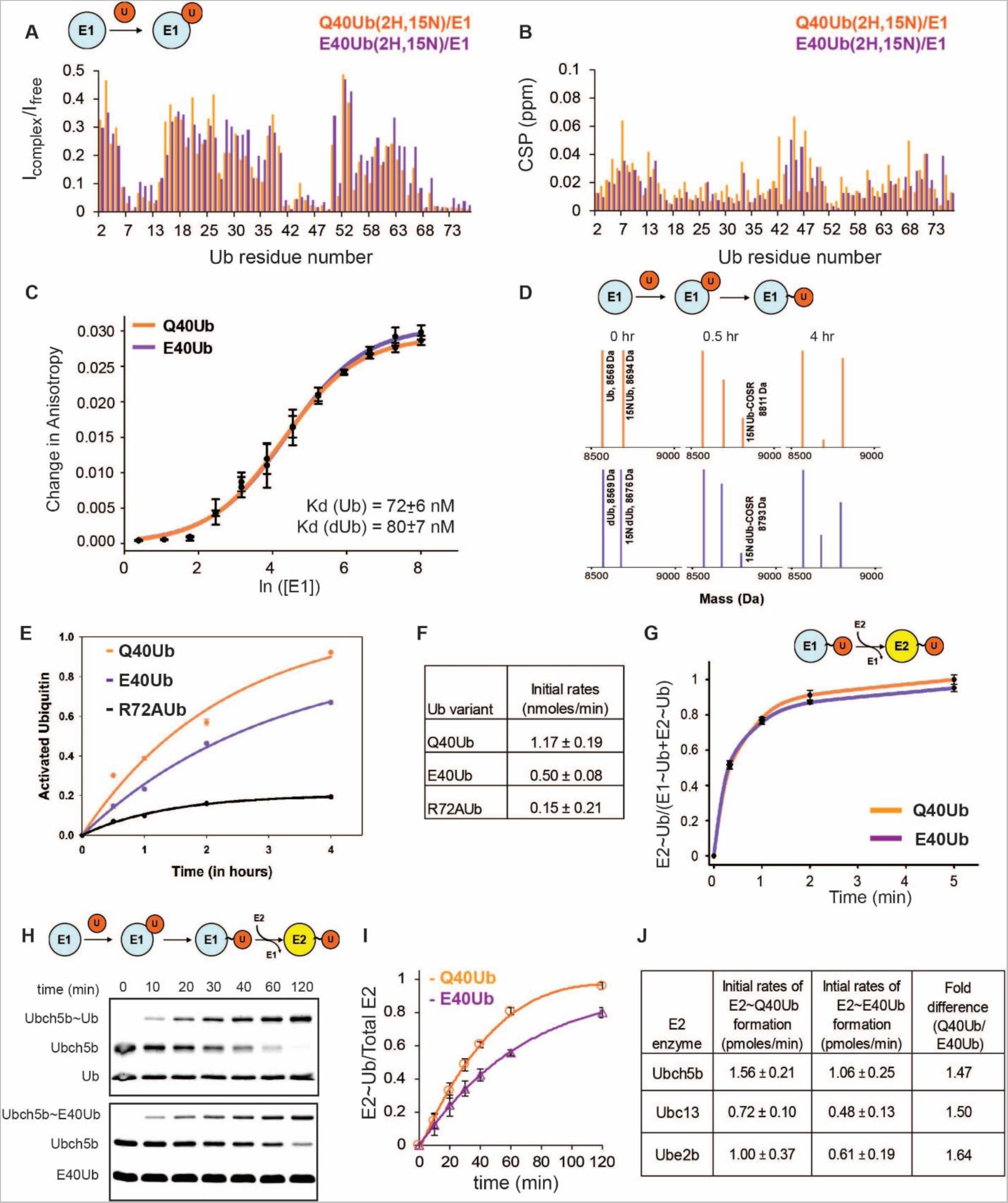
Deamidation retards the rate of E1 activation. (A) The ratio of Ub backbone amide resonance intensities (I_complex_/I_free_) in the Q40Ub/E1 and E40Ub/E1 non-covalent complex measured by ^15^N-TROSY-HSQC is plotted against the Ub residue numbers. The Ub molecules were isotopically labeled with ^2^H and ^15^N. I_complex_ is the intensity of Ub resonances in the Ub/E1 complex and I_free_ if that of free Ub. Extended data in Supp fig XX. (B) The CSPs in Ub in the Q40Ub/E1 or E40Ub/E1 complexes are plotted against Ub residue numbers. (C) Change in the fluorescence anisotropy of Q40Ub or E40Ub plotted against increasing E1 concentration is plotted, which yielded the dissociation constant of the E1/Ub complex. The Ub molecules were labeled with Alexa488. (D) ESI-MS analysis of the rate of E1~Ub conjugate formation in Q40Ub and E40Ub. Extended data in Fig S6. (E) The fraction of E1-conjugated Q40Ub and E40Ub is plotted against time. The R72A-Ub is plotted as a control. (F) Rate of E1~Ub to E2~Ub trans-thiolation is plotted as the fraction (E2~Ub)/((E1~Ub) +(E2~Ub)) against time. The Ub molecules were labeled with Alexa488. Extended data in Supp fig XX. (G) A plot of E2~Q40Ub and E2~E40Ub conjugation against time for the E2 UbcH5b. (J) The E2~Q40Ub and E2~E40Ub formation rates were measured for the E2s UbcH5b, Ubc13, and UbE2B.

After initial non-covalent E1/ubiquitin binding, the C-terminal tail of ubiquitin gets adenylated in the presence of Mg^2+^/ATP^18^. A chain of conformational reorientation in E1 culminates to form the thioester-linked E1~Ub^18, 19^. A Mass Spectrometry (MS) based method was used to measure the E1~Ub conjugate formation rate. If MESNa salt is added to the reaction mix, it rapidly attacks the covalently linked E1~Ub to form Ub-COSR. The rate of ubiquitin activation by E1 can be determined by quantifying Ub-COSR produced over time by MS. The rate of Q40Ub activation by E1 was 1.15 ± 0.19 nmoles/min (Figure 3D-F, S6C). The R72Aubiquitin mutant, known to abrogate E1 activation, acted as a negative control and had minimal activation. The rate of E1~E40Ub conjugation was 0.50 ± 0.08 nmoles/min, a two-fold reduction from the wild-type. R72/R74 has multiple interactions with E1 in the E1~Ub conjugate (Figure S7) and plays a crucial role in the formation of E1~Ub^21^. The intermolecular interactions of R72 and R74 with E1 will be perturbed by the intramolecular interactions with the D39/E40 acidic patch in E40Ub, reducing the rate of E1~Ub conjugation. Overall, deamidation does not impact the non-covalent E1/ubiquitin interaction but hampers the ubiquitin activation by E1.

E1 transfers the activated ubiquitin to the active site cysteine of the conjugating enzyme E2 via a trans-thiolation reaction. The transfer of Alexa FluorTM 488 tagged Q40Ub/E40Ub from E1 to Ubc13 (E2 enzyme) was measured to examine if deamidation could affect trans-thiolation. The trans-thiolation rate between Q40Ub and E40Ub had no detectable differences (Figure 3G, S8). When the E2~Ub conjugation rate was monitored in the presence of E1, Ub, ATP/MgCl_2_, the rate of E2~Ub conjugation was reduced in E40Ub by 1.5 fold (Figure 3H-J). The fold difference was conserved across different E2 families (UbcH5B, Ubc13, and UbE2B), suggesting that the effect of deamidation is not E2-specific. The difference in E2~Ub conjugation was similar to the difference in E1~Ub activation. Altogether, deamidation reduces the rate of E1~Ub activation, which is translated to a lower rate of E2~Ub formation in the ubiquitination reaction.

### The intramolecular salt bridge destabilizes the E2~Ub catalytic conformation

In the E2~Ub/E3 complex, Q40 is positioned at the ternary interface between E2, Ub, and E3. Deamidation may perturb the ternary interaction network and inhibit E2~Ub activation by E3s. Using the crystal structures of E2~Ub/RING complexes, we carried out an MD simulation of E2~Q40Ub/RING and E2~E40Ub/RING complexes. There was no discernible instability of ubiquitin in the closed E2~Ub/RING conformation in Q40Ub versus E40Ub (Figure S9), suggesting that the Q/E substitution does not impact once the ternary closed conformation is formed.

A few E2s, like Ubc13/Mms2 and E2-25K, have basal ubiquitination activity without E3s. We tested their activity with Q40Ub and E40Ub (Figure 4A, B). E40Ub inhibited the ubiquitination activity of these E2s in the absence of E3. The basal activity of these E2s suggests that they form the E2~Ub closed active conformation without E3. The E3s select this closed conformation to bind, stabilize and further activate it. Deamidation may impact the stability of the E2~Ub closed conformation (Figure 4C), which will affect its binding to RING and activity. Multiple Steered MD simulations (SMD) were performed wherein ubiquitin was subjected to forced dissociation from Ubc13. Differences in the average unbinding force/work between Q40Ub and E40Ub conjugates indicate the closed state’s destabilization. The unbinding force and work were reduced by ~96 pN and ~9 kcal/mol, respectively, for the E40Ub conjugate (Figure 4D, 4E), suggesting that Ubc13~E40Ub is unstable compared to Ubc13~Q40Ub. The interface of Ubc13 and ubiquitin is mediated through the I44 hydrophobic patch in ubiquitin. The reduced stability is similar to the I44A-Ub, where the I44 interactions were nullified. The stability was largely regained by substituting D39A or R72A, indicating that the deamidation-induced salt bridges destabilize the closed state. Interestingly, the effect was not reversed in the R74A mutant, suggesting that when ubiquitin C-terminal end is conjugated to E2 active site, R74 may be lodged at the active site cleft and may not efficiently form the intramolecular salt bridge with the D39/E40 patch.

**Figure 4.**
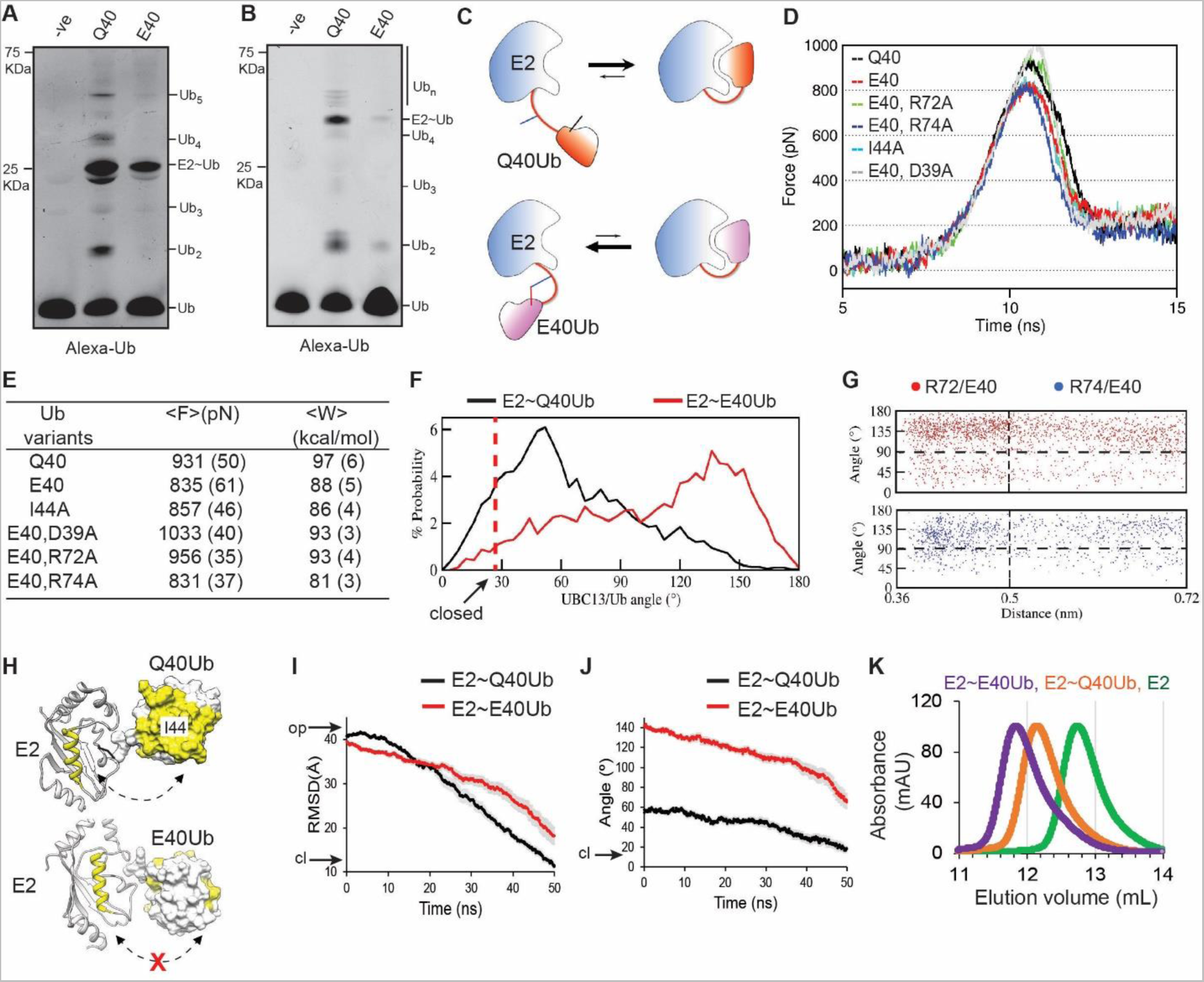
Deamidation perturbs the E2~Ub conformational equilibrium. (A) In-vitro ubiquitination reaction was performed using Ubc13 as the E2 and E40Ub or Q40Ub for 20 min. Mms2 was used as a co-factor in the reaction. The –ve lane is the same reaction without ATP. (B) Same as in (A) using Ube2K as the E2. (C) The model of deamidation induced perturbation of the E2~Ub dynamics. (D) Mean force-time profiles were obtained from Steered MD simulations of Ubc13~Ub and various Ub mutants. (E) Mean F_max_ and unbinding work (W) for the dissociation of Ub from the Ubc13~Ub closed-state. One standard error of the mean is indicated in the brackets. (F) Combined probability distribution of Ubc13/Ub intermolecular angle in the Ubc13~Q40Ub and Ubc13~E40Ub trajectories. (G) 2D plots showing the correlation between intramolecular salt-bridge and Ubc13/E40Ub angles. The higher values of angles correlate with the formation of E40 salt bridges. (H) Representative structures of high probability state in the E2~Q40Ub and E2~E40Ub conjugates. The hydrophobic patch in Ub and the central α-helix α2 are colored yellow. (I) The change in RMSD of Q40Ub (black) and E40Ub (red) is plotted against time from targeted MD simulations of E2~Ub from open to closed conformation. The RMSD is calculated relative to Ub in the E2~Ub closed conformation. (J) The angle between Ubc13 helix II and Ub helix I from the same trajectory is plotted against time. (K) The elution profile of E2 and E2~Ub in size-exclusion chromatography.

Multiple independent unbiased simulations (5 x 0.2 μsec) were carried out on the Ubc13~Q40Ub and Ubc13~E40Ub starting from the open state, where the folded region of ubiquitin (residues 1-71) do not interact with Ubc13. Interestingly, during the simulations, the folded ubiquitin region interacted with Ubc13 multiple times, suggesting that ubiquitin samples a conformational landscape in the open state (Figure S10 and S11). The new intramolecular salt bridges formed spontaneously during the simulation (Figure S12). To assess the possible effect of salt bridges on the conformational landscape of Ubc13~Ub, representative conformations obtained from conjugate ensembles were compared using RMSD-based structural clustering (Materials and Methods). Representative conformations of the first ten clusters were analyzed for both ensembles (Figure S13). An intriguing feature of the Ubc13~E40Ub ensemble was the inversion of E40Ub orientation for Ubc13. The ubiquitin orientations were analyzed across all trajectories by calculating the angle between two predefined vectors, one along the helix of ubiquitin and the other along helix-II of Ubc13. The angle probability distribution indicates that the Ubc13~E40Ub ensemble is biased toward a relative inversion of ubiquitin (Figure 4F). 2D plots of Ubc13/ubiquitin angle as a function of R72/74-E40 distance suggest that the inversion (Ubc13/ubiquitin angle >90°) is correlated with the formation of the new salt bridges (Figure 4G and S14). MD simulations of the closed and open states of Ubc13~Ub collectively indicate that the salt bridges formed upon deamidation can (i) enhance dissociation of the E2~Ub closed state and (ii) populate the E2~Ub open state conformations that are unfavorable for transition to the closed state (Figure 4H).

### Deamidation perturbs the open-to-closed transition in E2~Ub

Maintaining the proper orientation of ubiquitin is essential as it allows for complementary interactions between helix-II of Ubc13 and the I44 patch of ubiquitin to facilitate the transition from the open to the closed state. To study if the altered orientation in E40Ub perturbs the transition from the open to the closed state, Targeted Molecular Dynamic (TMD) simulations were performed. The starting structures were the open-state cluster representatives chosen from conventional MD simulations of Ubc13~Ub structures. The target structure was the closed-state crystal structure Ubc13~Ub (PDB: 5AIU). From the open state, Ubc13~Q40Ub moved towards the closed state, whereas the transition was retarded in Ubc13~E40Ub (Figure 4I, 4J). Hence, the Ubc13~E40Ub should have a higher population of open state conformations than Ubc13~Q40Ub.

The open state has a relatively larger dimension than the closed state. Isopeptide-linked Ubc13~Q40Ub/E40Ub conjugates (Figure S15) were produced, and a qualitative estimate of their dimensions was determined by analytical size exclusion chromatography. Proteins with larger dimensions elute earlier in the size exclusion chromatography. The Ubc13~E40Ub conjugate eluted earlier than the Ubc13~Q40Ub conjugate, confirming the higher population of extended open conformations in Ubc13~E40Ub. The analysis of the gel-filtration elution volumes yielded a Stokes radius of 24.5 Å and 23.1 Å for the Ubc13-E40Ub and Ubc13-Q40Ub conjugate, respectively. Upon deamidation, these experiments revealed a higher population of inactive, open E2~Ub conformation.

### The closed conformation of E40Ub is distinct from Q40Ub

The interactions between Ubc13 and E40Ub in the Ubc13~E40Ub conjugate were examined by NMR spectroscopy. The Ubc13~Q40Ub was purified, and CSPs were measured in both Ubc13 and ubiquitin compared to their respective free forms. High NMR CSPs were detected in the Ubc13 β2-β3 loop, the 3^10^ helix, helix α2 and α2α3 loop (Figure S16A). The loops and the 3^10^ helix surround the active site, and their CSPs suggest the ubiquitin conjugation at the active site. The α2 CSPs are reflective of the E2~Ub closed state. The hydrophobic patch (L7, V70, and I44), I36, and C-terminal end showed major CSPs in Q40Ub (Figure S16B). The C-terminal CSPs suggest conjugation to E2s, and the CSPs on the hydrophobic patch reflect the closed state. The CSPs are consistent with previous studies and the closed-state x-ray structure of Ubc13~Q40Ub. In the case of Ubc13~E40Ub, the CSP pattern is distinct from Q40Ub (Figure S16C, D). The major differences in CSPs were observed in the central helix II in Ubc13 (residues 100-110) (Figure 5A). In the Ub, the significant differences were around L8 and I44 hydrophobic regions (Figure 5B). The CSP differences suggest a lower population of the closed state and altered conformation of the E2~Ub closed state.

**Figure 5.**
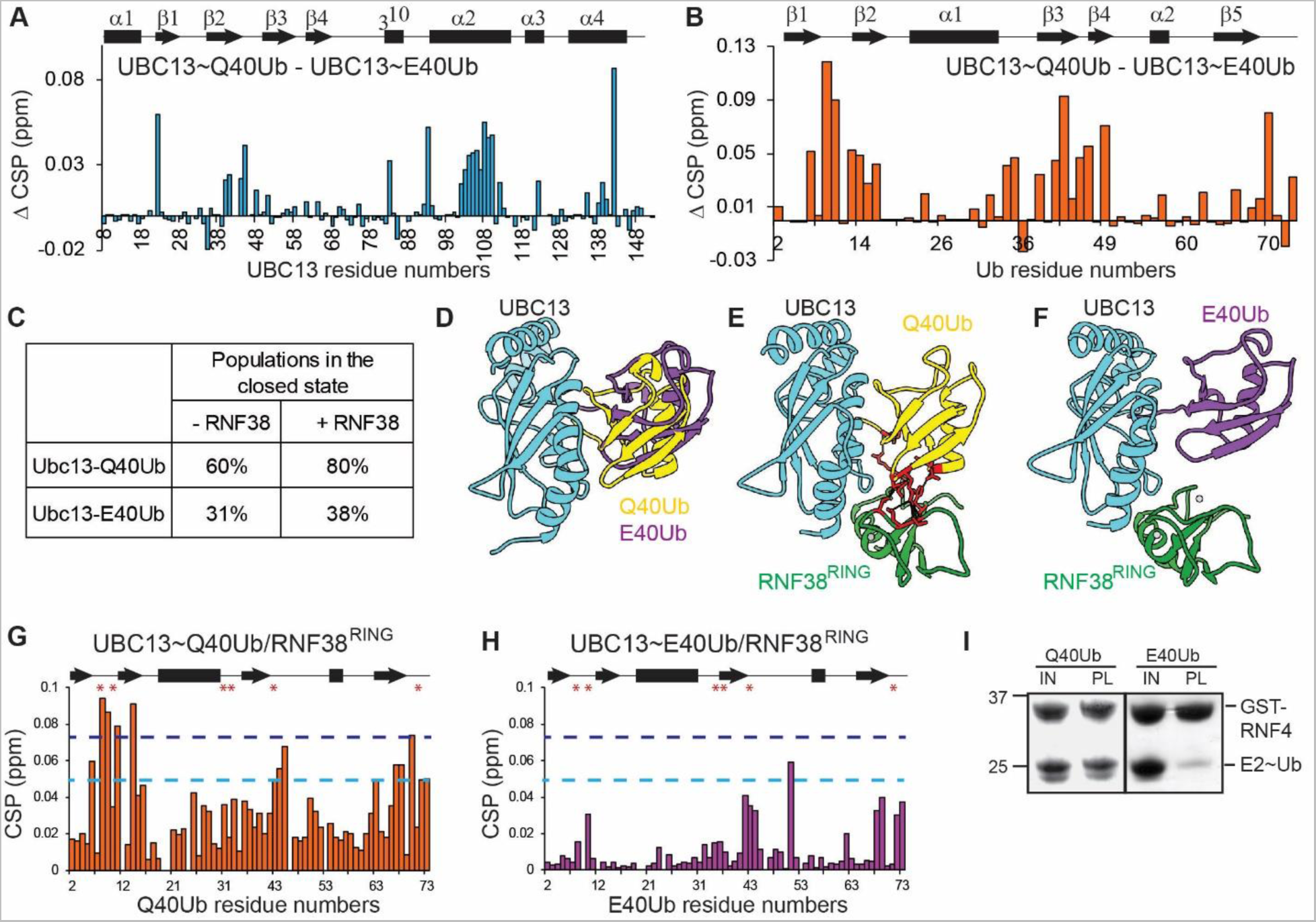
The distorted conformation of E2~E40Ub fails to bind RING domains. (A) The difference in CSPs (ΔCSPs) of Ubc13 when conjugated to Q40Ub and E40Ub. ΔCSP = CSP(Ubc13~Q40Ub) – CSP(Ubc13~E40Ub). (B) Same as in (A) is plotted for Ub. (C) The population of open and closed states calculated for the Ubc13~Ub/RNF38^RING^ complex is calculated by the model provided in Figure S16. (D) Ubc13~Q40Ub and Ubc13~E40Ub model structures generated using NMR CSPs by HADDOCK show differences in Ub orientation between the open and closed conformation. (E) A Ubc13~Q40Ub/RNF38^RING^ model suggests multiple interactions between Ub and RNF38^RING^ that stabilize the closed conformation. (F) The Ub/RING interactions are absent in a similar Ubc13~E40Ub/RNF38^RING^ model. (G) CSPs in Ub are plotted for the Ubc13~Q40Ub/RNF38^RING^ complex. The dark blue and light blue dashed lines correspond to Mean+2*SD. The residues in the UB/RNF38^RING^ interface observed in (D) are marked with asterisks. (H) CSPs in Ub are plotted for the Ubc13~E40Ub/RNF38^RING^ complex. The lack of high CSP values indicates a loss of interaction between E40Ub and the RING domain. (I) Pull-down assay with GST-RNF4^RING^ and Ubc13~Ub shows loss of RING binding upon deamidation.

To measure the changes in open and closed state populations due to deamidation, we used NMR spectroscopy. RING domains contact ubiquitin in the closed E2~Ub conformation but not the open conformation. Assuming that the affinity of RING for free E2 is similar to the affinity for E2~Ub open conformation (where ubiquitin does not contact RING), the equilibrium constant of open-to-close transition and the populations of open/closed states relate to the E2:RING and E2~Ub:RING dissociation constants (Figure S16E)^33^. We chose the RING domain from E3 RNF38 (RNF38^RING^) domain as the E3-RING for this study. ^15^N-labeled Ubc13 was titrated with RNF38^RING^, which yielded a K_d_ of 55 μM. Ubc13~Q40Ub and Ubc13~E40Ub were titrated with RNF38^RING^ to retrieve their K_d_ values (Figure S16F). Using the K_d_ values and the ratio of peak CSPs, the rate constant K and populations were calculated (details in Materials and Methods). The population of Ubc13~Q40Ub closed state was 60% in the absence of RNF38^RING^ and increased to 80% in the presence of RNF38^RING^ (Figure 5C). The population of the closed state was lower in Ubc13~E40Ub (30%), which meagrely increased to 38% in the presence of RNF38^RING^.

The Ubc13~Q40Ub and Ubc13~E40Ub complex was modeled in the HADDOCK using the NMR CSPs. The C-terminal end of ubiquitin was restrained to the active site of Ubc13 to mimic the native thioester conjugate. The Ubc13~Q40Ub complex was similar to the crystal structures of the closed state observed in previous studies^34^. The Ubc13~E40Ub complex showed a twisted orientation of E40Ub with respect to Q40Ub in the closed state (Figure 5D). The region around residue L8 is disoriented in E40Ub from the native conformation. Overall, deamidation modulates the active Ubc13~Ub conformational dynamics in two ways, i) reduce the open-to-closed state transition, ii) the disorient the ubiquitin in the closed state.

### RING-E3 fails to stabilize E2~Ub upon deamidation

RING forms several contacts with ubiquitin in the E2~Ub/RING closed complex. The distorted orientation E40Ub in the E2~Ub closed state may impact binding with RING. When RNF38^RING^ is modeled on the Ubc13~Ub haddock structures with E2~Ub/ RNF38^RING^ structures as a template, RNF38^RING^ forms several contacts with Q40Ub but not E40Ub (Figure 5E, 5F). The RING/ubiquitin contacts increase the affinity of RING for E2~Ub by several folds^35^. The effect of deamidation on RING/ubiquitin contacts was examined by measuring the CSPs on ubiquitin in the Ubc13~Ub conjugate upon the addition of RNF38^RING^ (Figure 5G). Residues in Q40Ub had high CSPs in the β1β2 turn, which is the binding interface of ubiquitin with RING. These CSPs were missing in E40Ub, suggesting a lack of interaction with the RING domain (Figure 5H). RNF38^RING^ functions as a monomer RING domain, whereas several RING-E3s, like RNF4^RING^ domains, function as homodimers^31^. The lack of RING/ubiquitin contacts was evident even in the case of a dimeric RING like RNF4^RING^ (Figure S17A). The region around L8 forms the majority of contacts with ubiquitin in the ternary complex. The disorientation around L8 in E40Ub breaks its contact with RING domains (Figure S17B). A pull-down assay was performed with GST-tagged RNF4 bound to GST beads and Ubc13-ubiquitin as the prey protein. The RNF4^RING^ could interact strongly with Ubc13~Q40Ub, but failed to bind the Ubc13-E40Ub conjugate (Figure 5I).

## Discussion

Bacterial deamidation has emerged as a powerful tool for regulating host signaling pathways. With a wide variety of target proteins like Rho GTPases, eukaryotic initiation factor 4A, UBLs (ubiquitin and NEDD8), and Ubc13, pathogenic bacteria employ subtle mechanisms to hijack immune response and evade detection^22^. Amongst an arsenal of effectors, glutamine deamidase catalyzes the irreversible conversion of specific glutamine residues of their target proteins into glutamate^15, 23^ and is a key virulence mechanism. *Legionella pneumophila* effector MavC catalyzes transglutamination between Ubc13 K92/94 and ubiquitin Q40, specifically inactivating Ubc13/MMS2 mediated K63 linked ubiquitin chains synthesis^24, 25^. *Shigella flexneri* employs effector OspI to deamidate Q100 of ubiquitin-conjugating enzyme Ubc13^23^, enabling E100/R14 intramolecular salt-bridge formation. The introduction of negative charge and salt-bridge competition perturbs both native and transient interactions of Ubc13 with RING E3s^26^. Enteropathogenic *Escherichia coli* secretes *cifs* to convert Q40 of NEDD8 to E40 that leads to the formation of E40/R74 intramolecular salt-bridge (Mohanty et al). Consequently, unfavorable interactions are promoted that inactivates Cullin RING Ligases (CRLs). *Cif*-homolog from *Burkholderia pseudomallei* (CHBP) targets a highly conserved Q40 in both ubiquitin and NEDD8 to bring cytopathic effects^15^. Ubiquitin deamidation drastically reduces the polyubiquitin chain synthesis in several E2/E3 pairs suggesting ubiquitin inactivation over any specific E2/E3 complex inactivation. Given the importance of the ubiquitin c-terminal tail in the ubiquitination reaction, any alteration in the tail could significantly affect its interactions and hamper broader sets of E2/E3 complexes.

Deamidation does not alter the globular ubiquitin structure. However, salt-bridge(s) between E40/D39-R72/74, observed in MD simulations and both NMR and crystal structure, restricts the flexibility of the c-terminal tail. The flexibility of the c-terminal tail is crucial for conjugation to E1 and E2 enzymes. Deamidation reduces the rate of ATP-dependent ubiquitin activation by E1 without affecting the E1/ubiquitin interaction or the trans-thiolation to E2s, lowering the cellular pool of available E2~Ub conjugates. Upon conjugation, the intramolecular salt bridges frequently invert the orientation of ubiquitin concerning E2, leading to an aberrant and reduced sampling of the total available conformational space compared to the wild-type conjugate. This orientation is incompatible with the conformational transitions required to achieve a catalytically closed state due to the hampered interactions between Ubc13 helix II and ubiquitin I44 hydrophobic patch. Our study provides detailed insight into mechanisms that disable E2~Ub conjugates to populate catalytically closed conformation upon deamidation.

E2/E3 interactions govern the downstream processes by specifying the substrate, linkage specificity, localization, etc. However, the lower sub-micromolar affinity range of E2/E3 interactions checks for any unregulated ubiquitination. E2~Ub conjugate enhances the RING affinity ~10-50 folds by facilitating more hydrophobic and electrostatic interactions that stabilize the dynamic E2~Ub ensemble to a catalytic conformation^26, 29, 30^. The E2~Ub/E3 ternary complex is stabilized by a highly conserved ‘Arg’ linchpin in RING E3s that makes hydrogen bonds with the side chains and backbone of both E2 and ubiquitin and restricts their orientation, allowing nucleophilic attack at the E2~Ub thioester^31^. NMR dynamics data reveals that the non-native conformations of E2~Ub upon deamidation limit interactions that stabilize ternary networks and reduce E3 binding. NEDD8 deamidation by cif effector inactivates cullin-RING ligases to check cell cycle progression and cytoskeletal regulation. Other deamidases like OspI and MavC silence immune responses. However, ubiquitin deamidation by CHBP inactivates E2~Ub, which could potentially hamper multiple signaling pathways downstream in the host cells. Thus, understanding the mechanism and consequences of E2~Ub inactivation has broader implications for host-pathogen interactions.

## Acknowledgments

The work was supported by intramural grants from the Tata Institute of Fundamental Research, Department of Atomic Energy, Government of India, under project no RTI 4006. The NMR data were acquired at the NCBS-TIFR NMR Facility, supported by the Department of Atomic Energy, Government of India, under project no RTI 4006. The X-ray crystallography data was collected at the X-ray facility at the NCBS Facility supported by the Department of Biotechnology, B-Life grant under project no dbt/pr12422/med/31/287/2014. The Mass spectrometry data was collected at the Mass-spectrometry facility at NCBS.

