## Supporting Information for "Bacterial deamidases modulate ubiquitin structure and dynamics to dysregulate ubiquitin signaling"

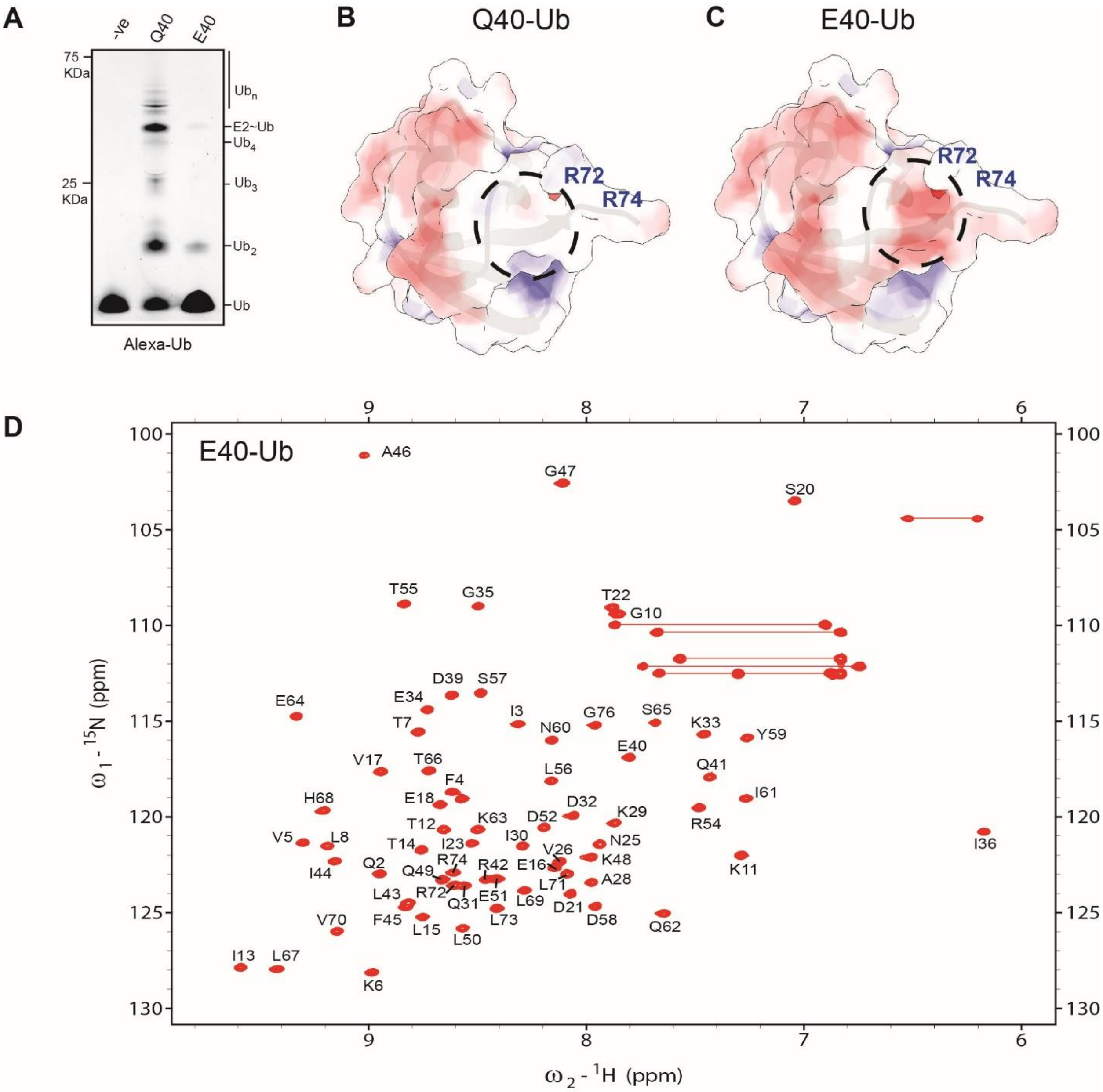

Figure S1. A) In-vitro ubiquitination reaction was performed using Ube2K as the E2, RNF38$^{RING}$ as the E3, and E40Ub or Q40Ub for 10 min. Mms2 was used as a co-factor in the reaction. The –ve lane is the same reaction without ATP. B) The surface electrostatic potential of Q40Ub was calculated through the Adaptive Poisson-Boltzmann Solver (APBS). The negatively charged surface is colored red, and the positively charged surface is colored blue. The color scale ranges from -8 to +8 KT/e. C) The same is plotted for E40Ub. The circled region highlights a negatively charged patch centered at E40. D) The $^{15}N$-$^{1}H$ HSQC spectra of E40Ub are plotted, and the backbone amide residues are indicated.

Table S1 Data-collection and data-processing statistics for the E40Ub. (Values in parentheses are for the highest resolution shell.)

| | |
|---|---|
| Space group | C121 |
| Temperature | 100K |
| X-ray source | Rigaku FR-X |
| Wavelength | 1.5148 |
| Resolution (Å) | 35.24 - 1.6 |
| Unit cell parameter | a = 48.01 Å, b = 39.06 Å, c = 35.29 Å, α = γ = 90.0°, β = 93.12° |
| Molecules per asymmetric unit | 1 |
| Matthews Coefficient ($Å^3$ $Da^{-1}$) | 1.94 |
| Solvent content (%) | 36.52 |
| Total no. of reflection | 60413 (2123) |
| No. of unique reflection | 8673 (415) |
| Multiplicity | 7.0 (5.1) |
| Completeness (%) | 99.5 (93.3) |
| Average I/σ(I) | 32.8 (16.9) |
| Rmerge[a] (%) | 3.4 (7.6) |
| **Refinement Statistics** | |
| Rfactor[b] (%) | 13.61 |
| Rfree[c] (%) | 18.15 |
| RMS bond length (Å) | 0.0269 |
| RMS bond angle (°) | 2.6575 |
| Overall B (Isotropic) from Wilson Plot | 13.924 |
| **Ramachandran Map** | |
| Most favorable region (%) | 100 |
| Additional allowed region (%) | 0 |
| Outlier region (%) | 0 |

[a] $R_{merge} = \sum_{hkl} \sum_{i=1} /I_i\,(hkl) - \bar{I}(hkl) / \sum_{hkl} \sum_{i=1} I_i\,(hkl)$

[b] $R_{factor} = \sum | F_o - F_c | / \sum F_o$

[c] $R_{free} = \sum | F_o - F_c | / \sum F_o$ *where the F values are test set amplitudes (5%) not used in refinement.*

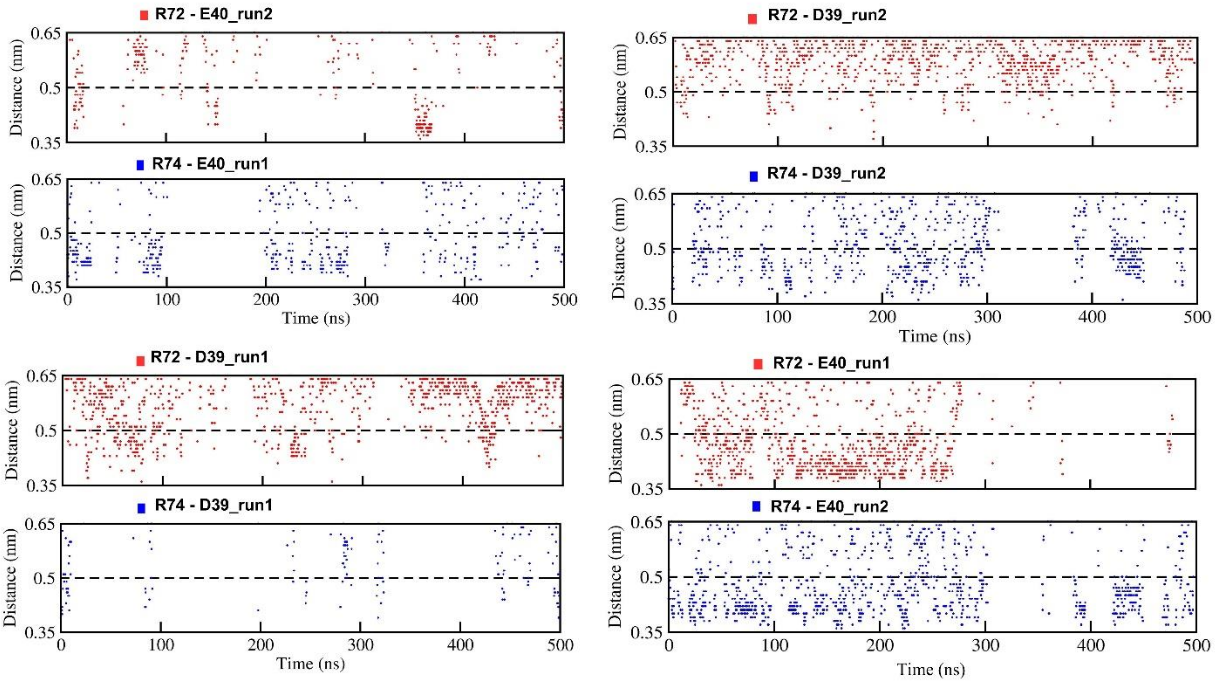

Figure S2. The distance between R72/R74-Cζ and E40-Cδ atoms or D39-Cγ is plotted for both the all-atom MD simulation runs. A distance of 0.5 nm (dashed line) was used as a cutoff to calculate salt bridge occupancies.

Table S2. NMR and refinement statistics of the p62UBA/Ub complex.

| | |
|---|---|
| Restraints | |
| Ambiguous restraints | 5 |
| Unambiguous restraints (Intermolecular NOEs) | 33 |
| Haddock Parameters | |
| Number of Clusters | 1 |
| Cluster Size | 200 |
| Haddock score | -72.7 (± 0.9) |
| Van Der Waals Energy | -31.6 (± 2.1) |
| Electrostatic Energy | -233.4 (± 9.9) |
| Restraints Violation Energy | +1.1 (± 0.3) |
| Buried surface area | +915.7 (± 27.0) |
| Rmsd (Å) | |
| All backbone | 0.5 |
| All heavy atoms | 0.6 |
| RMS deviation | |
| Bond Angles | 0.5° |
| Bond lengths | 0.004Å |
| Molprobity[b] | |
| Clash score | 7.9 |
| Score | 2.2 |
| Cβ deviations > 0.25Å | 0.0% |
| Bad Angles | 0.0% |
| Bad Bonds | 0.0% |
| Ramachandran Map | |
| Most Favoured | 98.9% |
| Allowed Regions | 1.1% |
| Disallowed Regions | 0.0% |

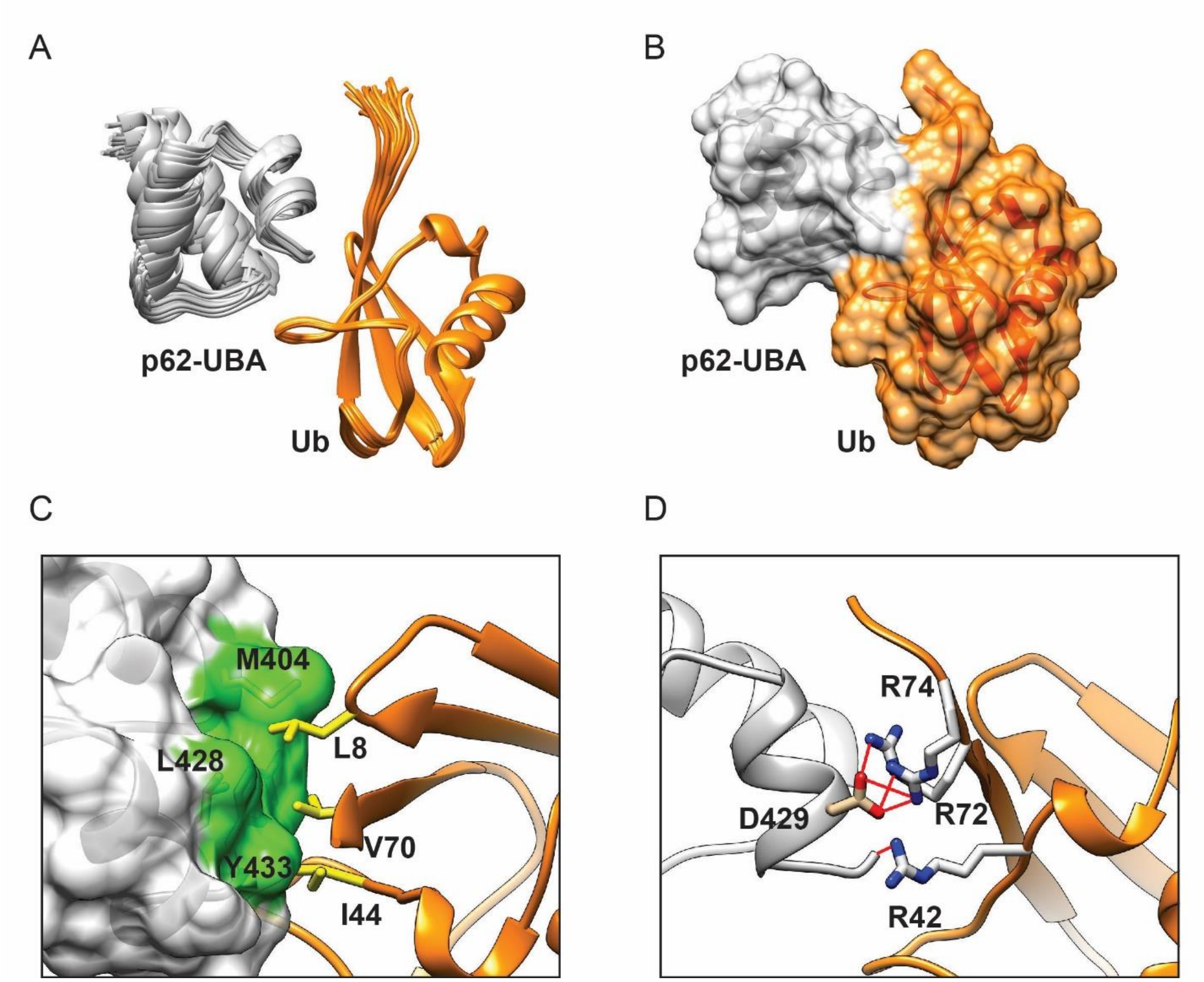

Figure S3. A) Twenty lowest energy structures of the p62-UBA/Ub complex, calculated by HADDOCK. p62-UBA is colored grey, and Ub is colored orange. The structures superimpose with low rmsd, suggesting that the calculation has converged. B) The surface representation of the lowest energy structure suggests an extensive interface between the UBA and Ub. C) The hydrophobic interactions at the p62-UBA/Ub interface are highlighted. The p62-UBA domain is shown as a surface where the hydrophobic regions are colored green, and the hydrophobic residues M404, L428, and Y433 are labeled. The sidechain atoms of Ubiquitin hydrophobic L8-I44-V70 are shown and labeled. D) The hydrogen bonds across the p62-UBA/Ub interface are shown. R72 and R74 form hydrogen bonds with D429. R42 contacts the backbone oxygen atom of Y433 to form a hydrogen bond.

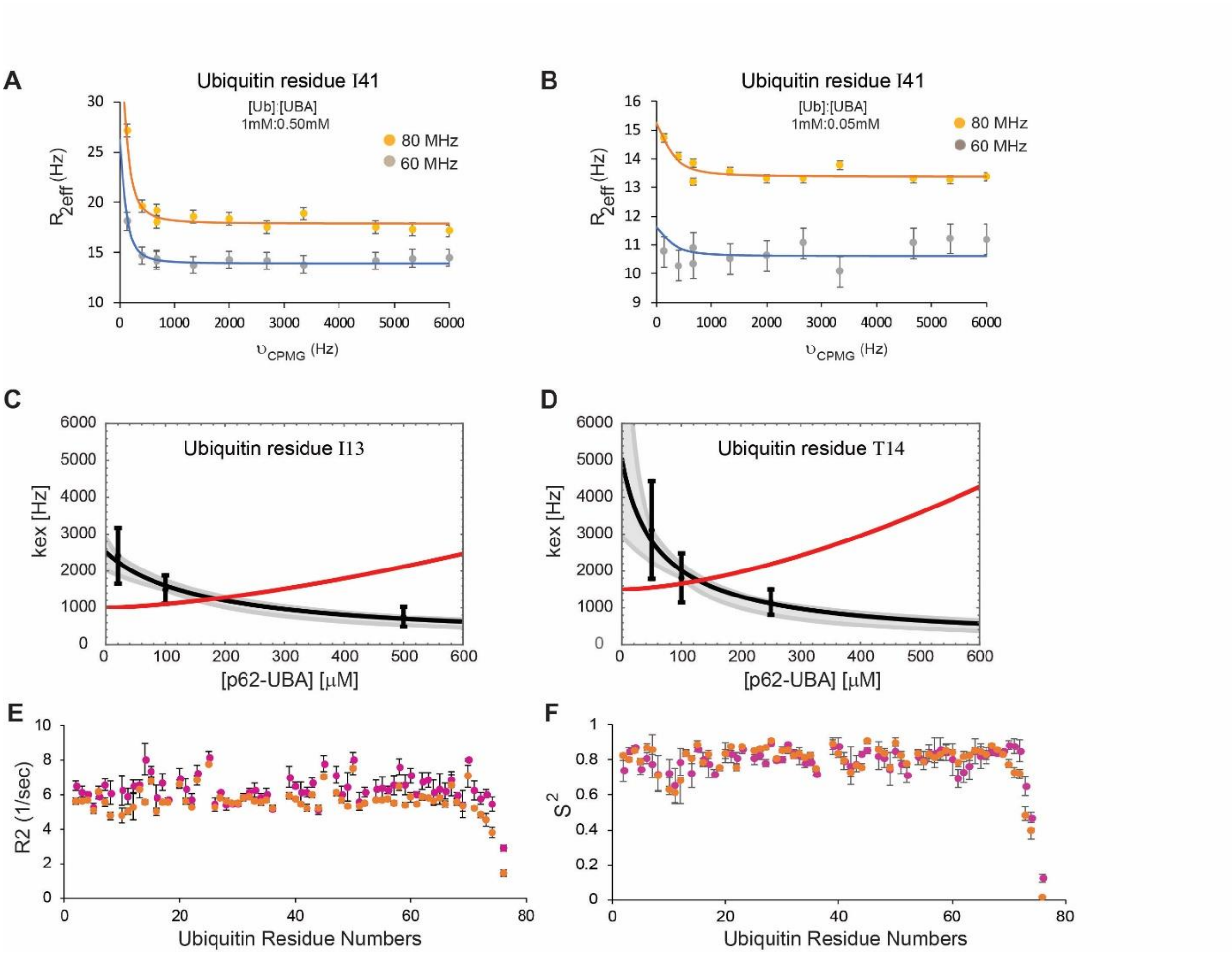

Figure S4. (A) $^{15}$N relaxation dispersion for the amide of the Q40Ub residue I41 at 80 MHz and 60 MHz in the presence of 0.5 mM of p62-UBA domain. (B) Same as (C) plotted for 0.05 mM of the p62-UBA domain. (C) and (D) The obtained exchange rates ($k_{ex}$) of the ubiquitin residue I13 and T14 (black data points with error bars) decrease with increasing p62-UBA concentration, which indicates conformational selection. The gray lines with shaded error regions result from fits of the ($k_{ex}$) equations of the two-state and conformational-selection binding mechanism. The red line denotes the induced fit model. E) and F) shows the $R_2$ and $S^2$ values for the Q40Ub (orange) and E40Ub (magenta) for all residues.

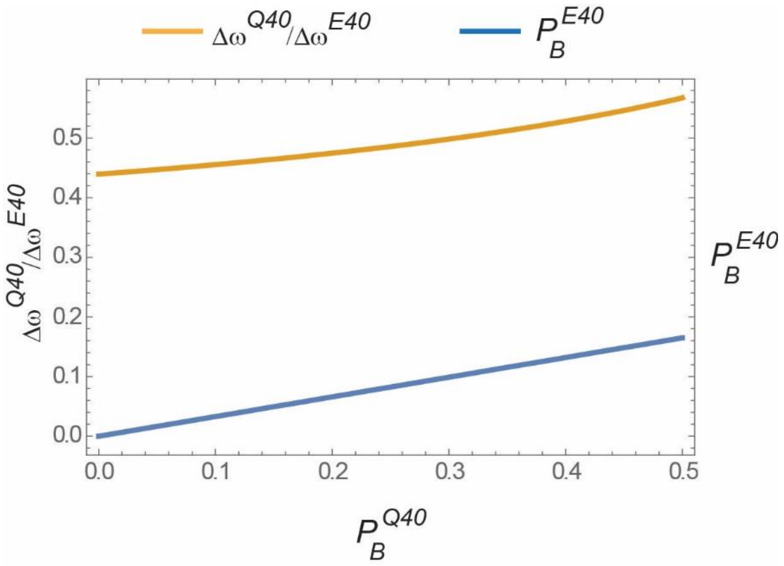

Figure S5. A plot of the population of a minor state in E40Ub ($P_B^{E40}$) against the population of a minor state in Q40Ub ($P_B^{E40}$) is shown as a blue line. The ratio of chemical shift differences between major and minor state in Q40Ub and E40Ub ($\Delta\omega^{Q40}/\Delta\omega^{E40}$) is plotted as an orange line.

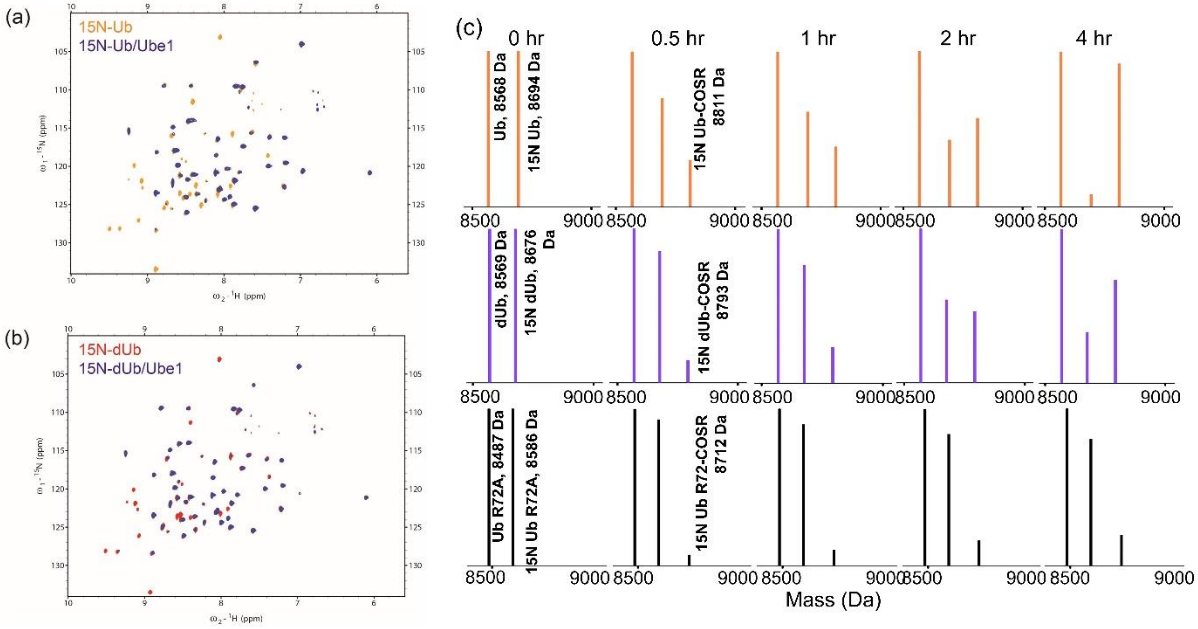

Figure S6. (a) Overlay of the 15N-edited HSQC spectra of free Q40Ub (orange) and in 1:1.5 complex with E1 (Purple). Identical spectra with E40Ub in (b). (c) ESI-MS spectral region of activated 15N-Q40Ub/E40Ub. N15Q40Ub/E40Ub incubated with E1, MESNa, ATP, and MgCl2 for indicated time points. Reaction quenched with EDTA and unlabelled Q40Ub added as the internal control. N15-Q40Ub/E40Ub-COSR peak can be seen increasing with time. R72AUb was used as a negative control.

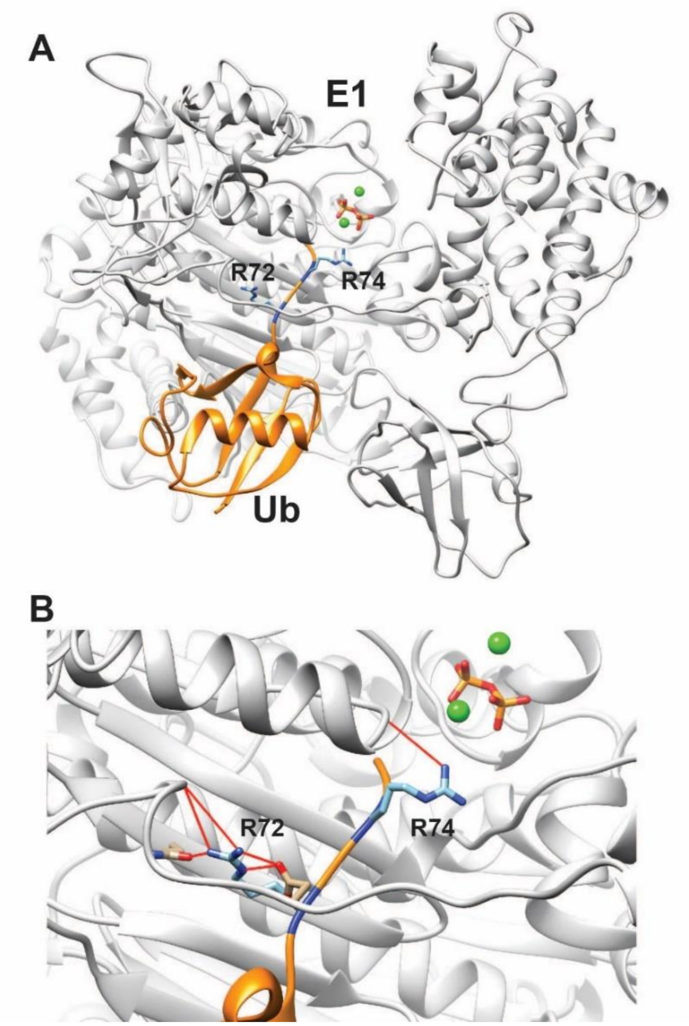

Figure S7. A) The structure of E1~Ub conjugate (PDB ID: 6CD6) is shown, where E1 is colored gray, and Ub is colored orange. The Mg2+ is colored green and the ATP analogue is shown. The sidechain of R72 and R74 are shown and colored blue. B) Hydrogen bonds between R72/R74 and E1 are highlighted as red lines.

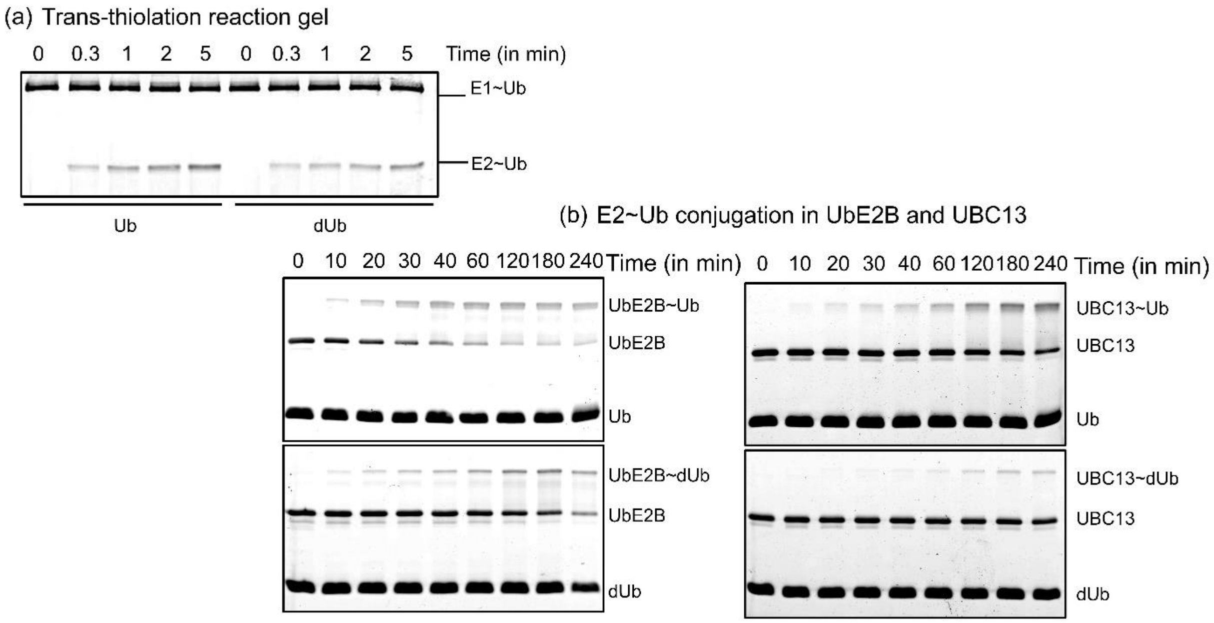

Figure S8. (a) Trans-thiolation was measured with Q40Ub and E40Ub discharge from E1~Ub with time and was run on 15% SDS-gel. (b) The rate of E2~Q40Ub and E2~~E40Ub conjugation was measured for the E2s Ube2b and Q40Ubc13. Active site cysteine to serine mutant used for charging. Reaction quenched in different time points and run on 15%SDS-gel and stained with SYPRO ruby red.

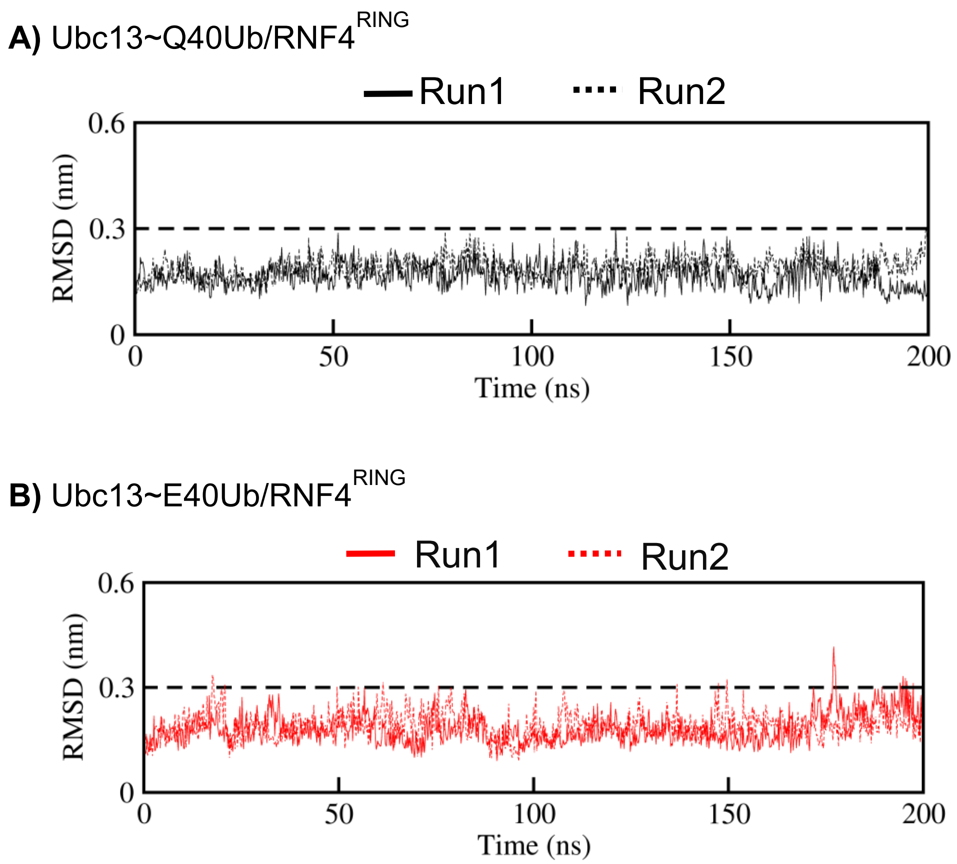

Figure S9. (A) The RMSD of Q40Ub in the Ubc13~Q40Ub/RING complex is plotted against time. Unbiased simulations were carried out from the X-ray crystal structure of Ubc13~Ub/RNF4$^{RING}$ complex (pdb id: 5AIU), where Ubc13~Ub is in the closed conformation. (B) The RMSD of E40Ub in the same complex is plotted against time. The Q40 was substituted to E40 in the same structure (pdb id: 5AIU) and simulated.

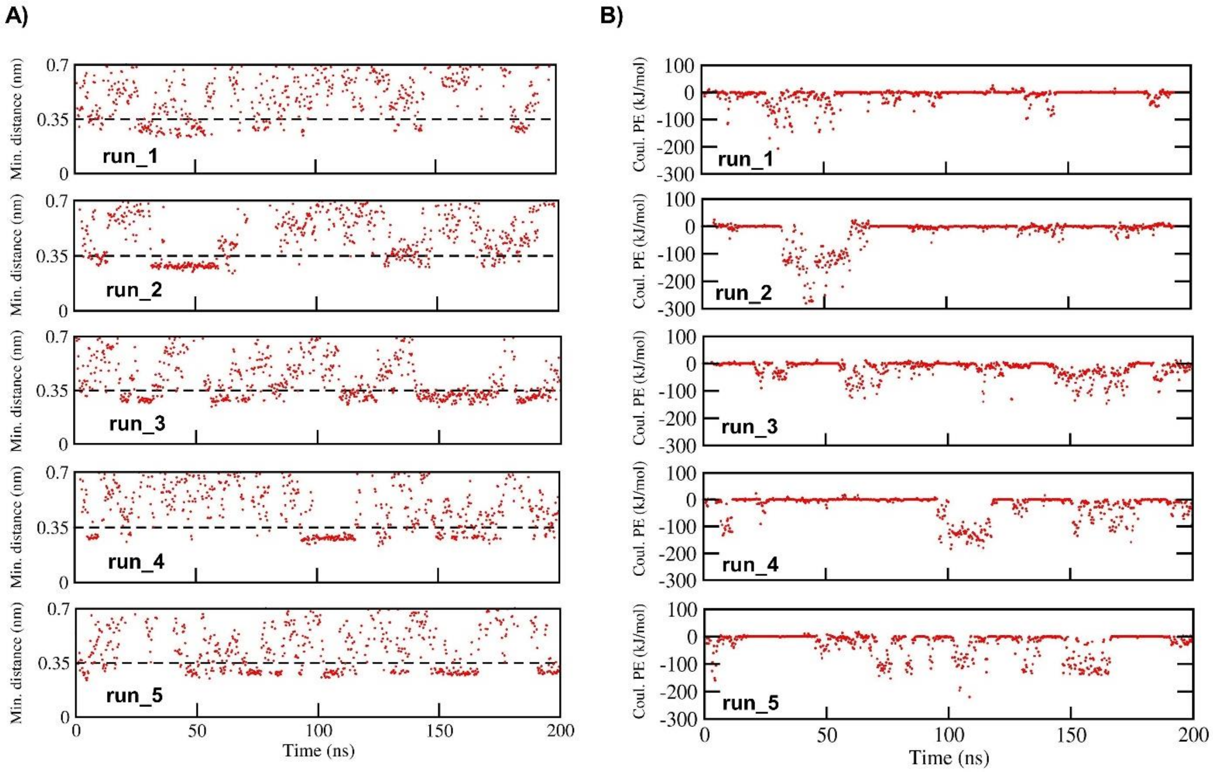

Figure S10. Dynamics and energetics of Ubc13~Q40Ub in the open state. A) The minimum distance versus time plots between Ubc13 and Q40Ub (residues 1-71) from five MD simulations (run_1 - run_5) initiated from the open state of Ubc13~Ub. A drop in the minimum distance value below 0.35 nm indicates one or more close-range interactions. B) Coulombic potential energy versus time plots between Ubc13 and Q40Ub (residues 1-71) for the same set of simulations as in A.

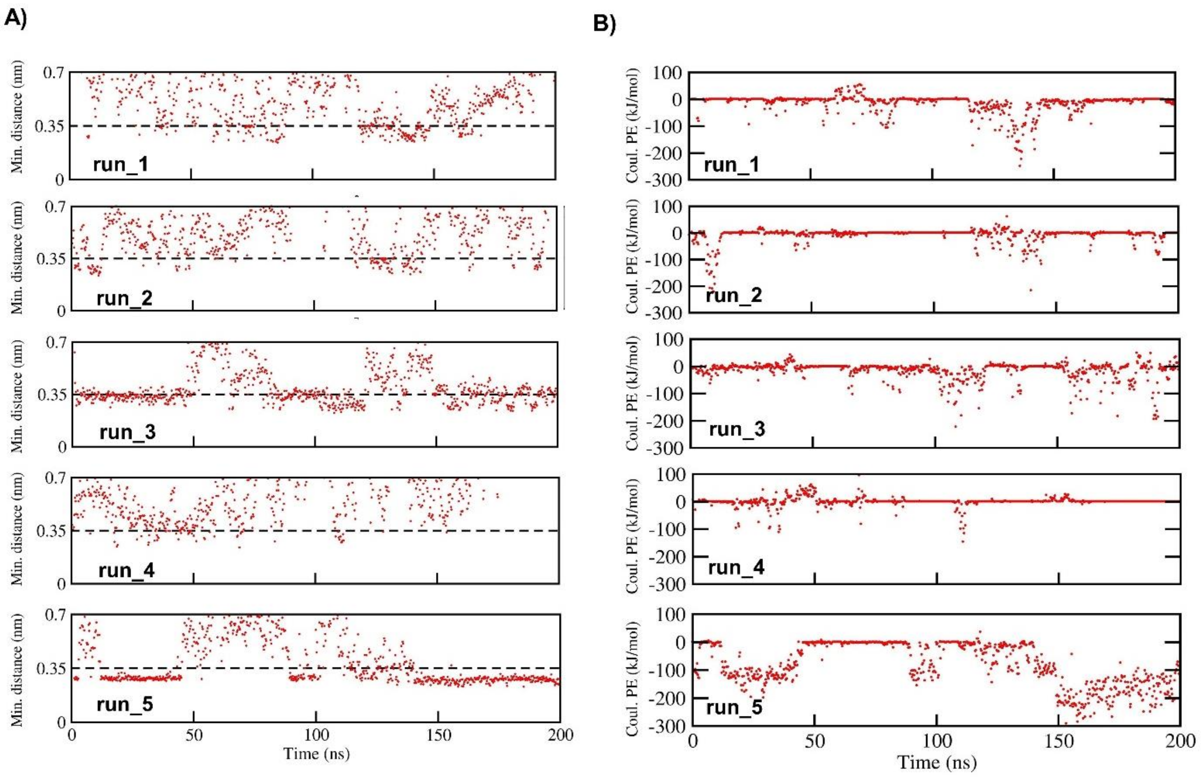

Figure S11. Dynamics and energetics of Ubc13~E40Ub in the open state. A) The minimum distance versus time plots between Ubc13 and E40Ub (residues 1-71) from five MD simulations (run_1 - run_5) initiated from the open state of Ubc13~Ub. A drop in the minimum distance value below 0.35 nm indicates one or more close-range interactions. B) Coulombic potential energy versus time plots between Ubc13 and E40Ub for the same set of simulations as in A.

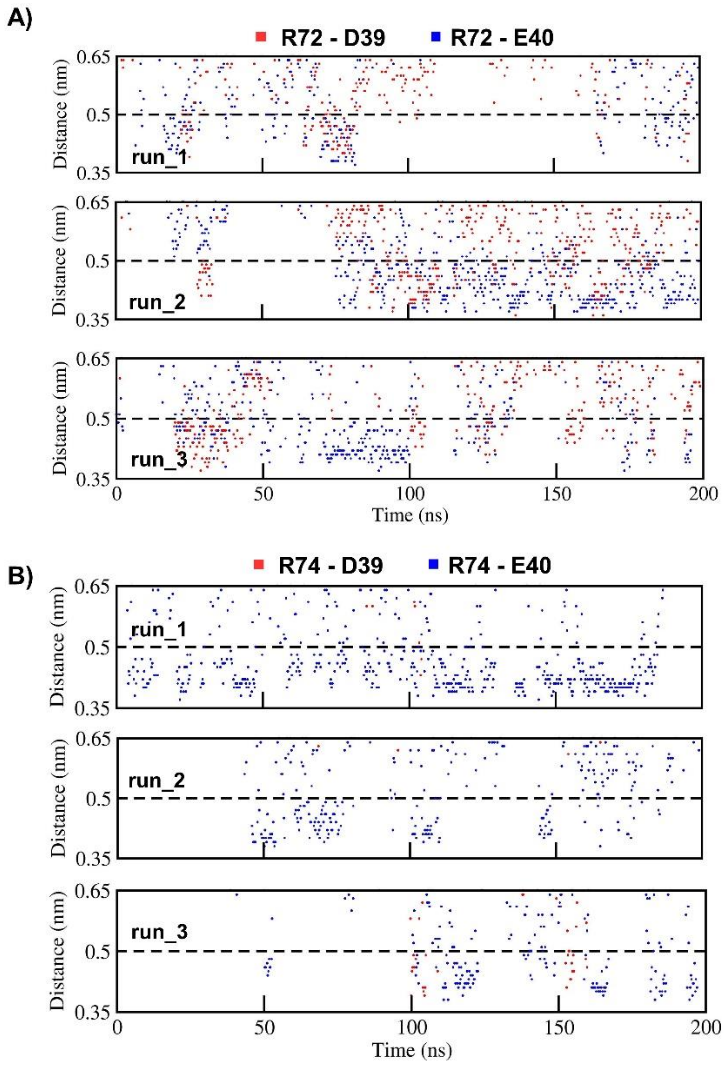

Figure S12. Dynamics of intramolecular salt bridges in E40Ub from MD simulations of Ubc13~E40Ub in the open state. Variation in salt bridge stability as a function of time across three independent trajectories for A) R72-D39 and R72-E40 salt bridge, and B) R74-D39 and R74-E40 salt bridge.

| Cluster Number | % of the total ensemble | |
|---|---|---|
| | Q40Ub | E40Ub |
| 1 | 15.4 | 12.8 |
| 2 | 8.8 | 11.5 |
| 3 | 8.3 | 7.1 |
| 4 | 7.5 | 6.6 |
| 5 | 7.1 | 6.1 |
| 6 | 5.3 | 6.1 |
| 7 | 5.0 | 5.1 |
| 8 | 4.1 | 4.9 |
| 9 | 4.0 | 3.9 |
| 10 | 3.1 | 3.1 |
| Total | 68.6 | 67.2 |

Figure S13. Percentage of the total ensemble for each of the ten most populated clusters from Ubc13~Q40Ub and Ubc13~E40Ub macrotrajectories.

A

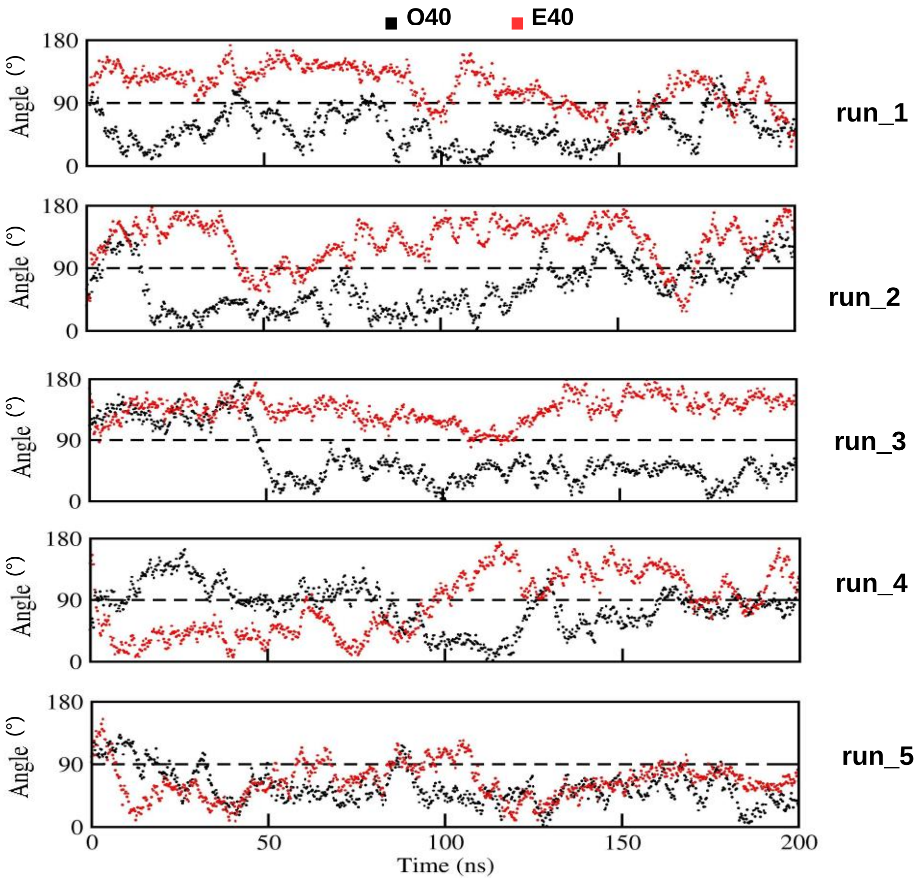

B

| Trajectory No. | Q40 | E40 |
|---|---|---|
| 1 | 52.7 (25.5) | 113.2 (28.6) |
| 2 | 62.1 (35.3) | 128.9 (30.7) |
| 3 | 62.9 (40.1) | 134.7 (19.2) |
| 4 | 80.4 (32.3) | 83.4 (44.8) |
| 5 | 56.3 (24.9) | 65.6 (25.2) |

Figure S14. (A) Ubc13/Ub angle variation as a function of time across five MD trajectories initiated from the open state for Ubc13~Q40Ub and Ubc13~E40Ub conjugates. (B) Mean Ubc13/Ub angle in degrees calculated from Ubc13~Q40/E40Ub trajectories.

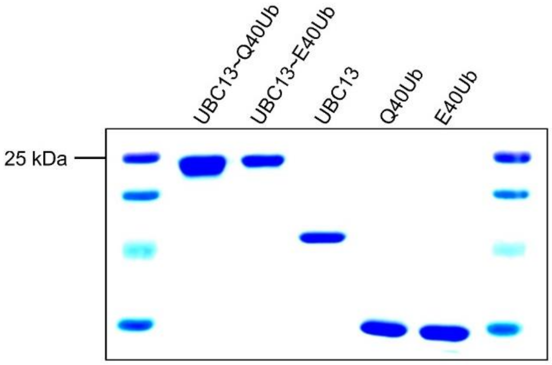

Figure S15. The SDS gel image of purified Ubc13~Q40Ub and Ubc13~E40Ub conjugates.

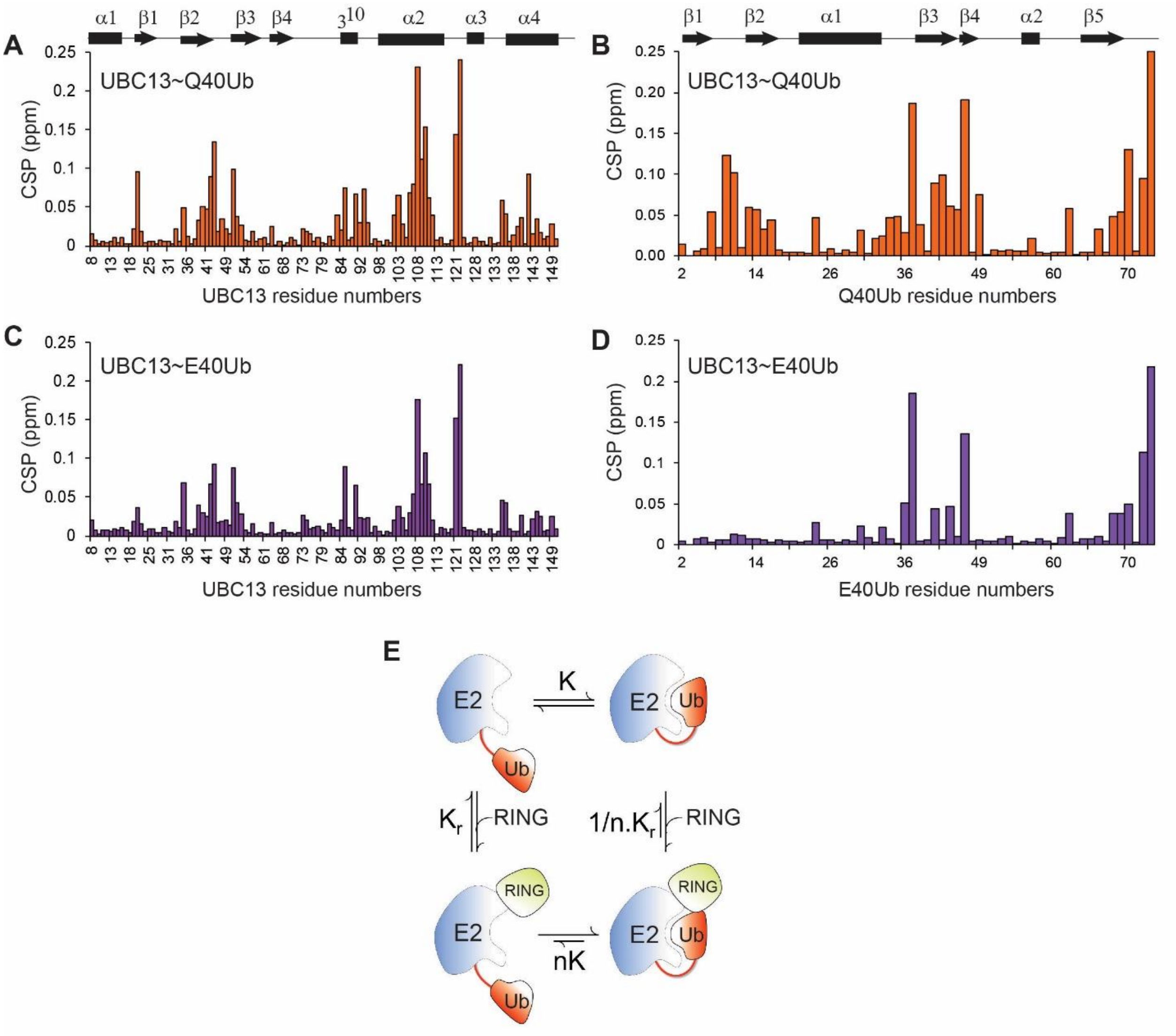

F

| | UBC13/RNF38 | UBC13~Q40Ub/RNF38 | UBC13~E40Ub/RNF38 |
|---|---|---|---|
| Dissociation constant (Kd) | 54.7 ( ± 2.7) | 19.6 ( ± 1.4) | 36.8 ( ± 2.2) |

G

| | | | K | |
|---|---|---|---|---|
| | n | r | - RNF38 | + RNF38 |
| Q40Ub | 2.57 ( ± 0.2) | 1.32 | 1.52 | 3.91 |
| E40Ub | 1.39 ( ± 0.2) | 1.24 | 0.45 | 0.63 |

Figure S16. The Chemical Shift Perturbations (CSPs) in A) Ubc13 and B) Ub in the Ubc13~Q40Ub complex is plotted against their residue numbers. The same is plotted for Ubc13~E40Ub in C) and D). E) The model of Ubc13 activation by the RNF38$^{RING}$ domain. F) The dissociation constants of Ubc13/RNF38$^{RING}$, Ubc13~Q40/RNF38$^{RING}$ and Ubc13~E40Ub/RNF38$^{RING}$ complexes. G) The values of n, r and K in Ubc13~Q40Ub and

Ubc13~E40Ub, determined from measured dissociation constants and CSPs. n is the change in rate of open-to-close conformational dynamics due to RING, and is calculated as the ratio of dissociation constants measured in presence and absence of Ub. r is the ratio of CSPs in Ubc13~Ub is the presence and absence of RING. K is the equilibrium constant of open-to-close motion and is calculated as $K=(n-r)/(nr-n)$. The populations calculated from K are plotted in Figure 5C.

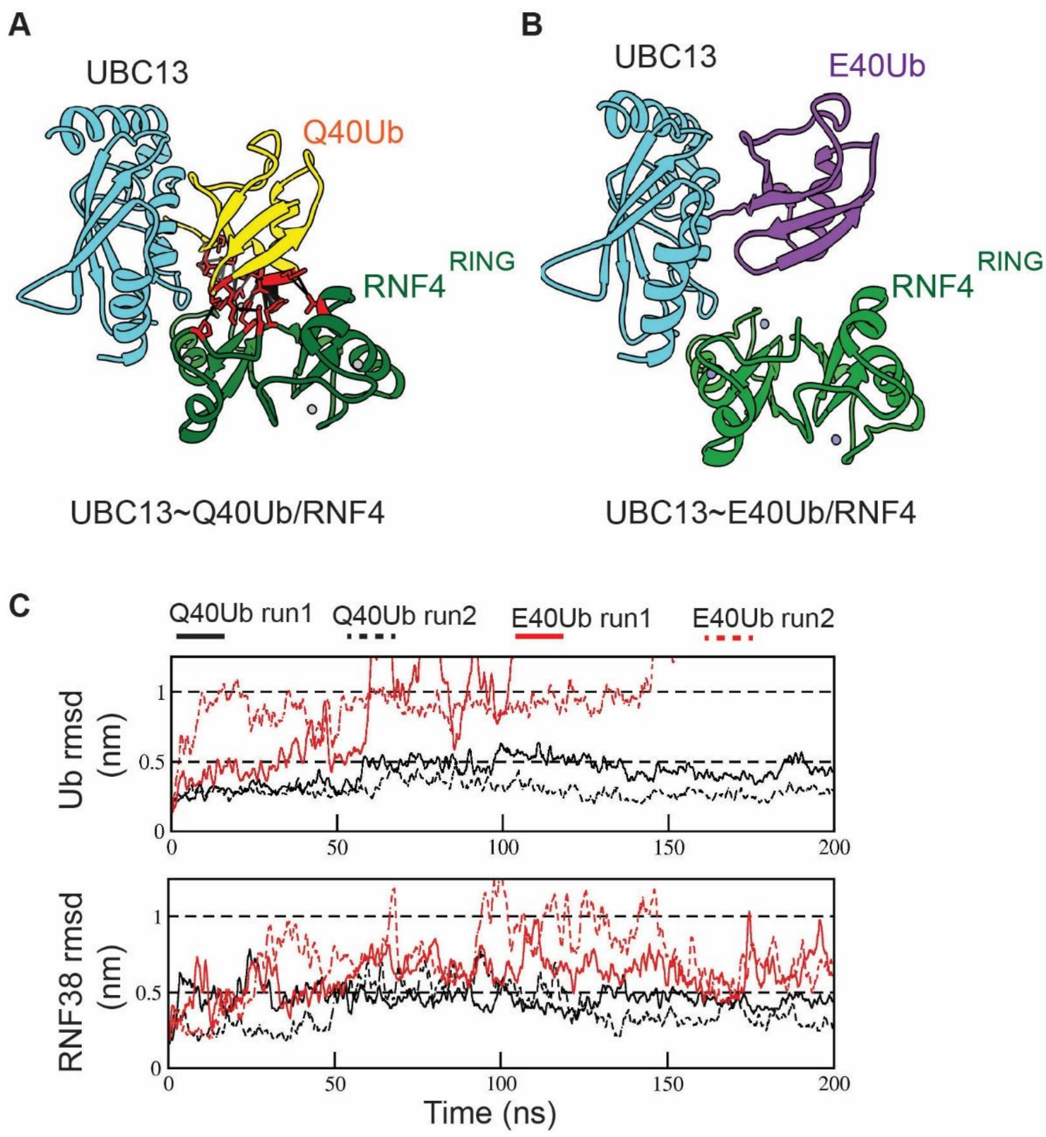

Figure S17. A) A Ubc13~Q40Ub/RNF4$^{RING}$ model suggests multiple interactions between Ub and RNF4$^{RING}$ that stabilize the closed conformation. (B) The Ub/RING interactions are absent in a similar Ubc13~E40Ub/RNF4$^{RING}$ model. C) The RMSD of Q40Ub and E40Ub in the ternary complex is plotted against time. Unbiased simulations were carried out from the structures given in Figure 5D) and 5E) for Q40Ub and E40Ub, respectively. The RMSD of RNF38$^{RING}$ is the same simulations are plotted in the below panel.
